# Persistent neuroimmune alterations in children who are HIV-exposed but uninfected at age 6–7 years: Associations with language development in a South African birth cohort

**DOI:** 10.64898/2026.03.03.709328

**Authors:** Cesc Bertran-Cobo, Frances C Robertson, Simone Williams, Tusekile Sarah Kangwa, Jenna Annandale, Jessica Ringshaw, Layla Bradford, Nadia Hoffman, Heather J Zar, Dan J Stein, Kirsten A Donald, Petrus J W Naudé, Catherine J Wedderburn

## Abstract

**Background:** Children who are HIV-exposed but uninfected (HEU) are at increased risk of neurodevelopmental delays, yet neuroimmune pathways linking perinatal HIV exposure to school readiness remain unclear.

**Methods:** In the Drakenstein Child Health Study, 268 children (94 HEU, 174 HIV-unexposed [HU]) underwent magnetic resonance spectroscopy in midline parietal grey and left parietal white matter regions at 6–7 years. Peripheral blood serum immune markers were measured in pregnancy and in children at 6 weeks, 2, 3, and 5 years. Linear mixed-effects models characterised child immune trajectories and linear regressions tested associations with creatine-referenced neurometabolite ratios and school readiness scores.

**Results:** Mothers living with HIV had higher sCD14 and lower MMP-9, NGAL, and GM-CSF than mothers without HIV (p<0.05). Perinatal HIV exposure was associated with altered trajectories of child sCD14, GM-CSF, IL-1β, IL-5, IL-10, and YKL-40. At 6–7 years, children who were HEU had lower parietal grey matter glutamate ratios and lower left parietal white matter choline ratios. By school entry, immune–neurometabolite associations were predominantly driven by child serum markers; IL-8 emerged as a consistent correlate across developmental stages. Children who were HEU had lower language scores than HU peers. Left parietal white matter choline ratios were positively associated with language and overall school readiness in HU children, but not HEU.

**Conclusions:** Perinatal HIV exposure was associated with alterations in immune development, neurometabolites reflecting both white matter maturation and neuronal health, and school readiness. Our findings highlight potential neuroimmune pathways contributing to neurodevelopmental risks in children who are HEU.

## Introduction

Globally, approximately 16 million children have been born to women living with HIV but remain uninfected themselves (1). Compared to their HIV-unexposed (HU) peers, children who are HIV-exposed uninfected (HEU) experience higher rates of mortality and morbidity (2,3), growth faltering (4), and neurodevelopmental delays (5), although these risks are not universal. Recent research prioritisation efforts have highlighted the need to identify which children are most vulnerable, and to uncover the pathophysiological pathways through which adverse outcomes may arise (6). This requires moving beyond descriptive comparisons toward longitudinal studies that can link early-life exposures to long-term neurodevelopmental outcomes.

Maternal immune activation during pregnancy is a biological mechanism that may contribute to neurodevelopmental risk in children. Studies have shown that dysregulated immune signalling *in utero* can disrupt neurogenesis, synaptic pruning, and myelination, with long-lasting consequences for cognition and behaviour (7–9). Pro-inflammatory cytokines have been found to cross the placenta and affect glial integrity and white matter maturation of the developing brain (7–11). Maternal infections have also been variably associated with changes in infant brain growth and connectivity (12). Among women living with HIV, chronic immune dysregulation is commonly documented and may persist despite antiretroviral therapy (ART) (13,14). This may result in priming of the developing brain, leading to exacerbated neuroimmune responses to postnatal risk factors (15,16).

In children who are HEU, maternal immune activation during pregnancy has been directly associated with altered immune (17,18) and cognitive development (19,20). However, findings on circulating cytokine and immune marker profiles are heterogeneous across settings and developmental stages: some studies report higher pro-inflammatory cytokines and monocyte activation markers in mothers living with HIV (21,22) and their children who are HEU (23,24), whereas others describe lower pro- and anti-inflammatory cytokine profiles (19,25), or no detectable differences between HEU and HU groups (26,27). Inconsistencies across studies may reflect socioeconomic and geographical differences (28), as well as participant’s age at the time of sampling, and specimen processing (29). With respect to neurodevelopment, data from a South African birth cohort, the Drakenstein Child Health Study (DCHS), showed that elevated cytokine levels in infants who are HEU at 6 weeks predicted poorer language and motor outcomes at 2 years (19). Similarly, a Kenyan study identified distinct immunologic predictors of neurodevelopmental outcomes based on perinatal HIV exposure (20).

Neuroimaging studies provide a key opportunity to explore the impact of maternal HIV status on child brain development; however, magnetic resonance imaging (MRI) data from areas of high HIV prevalence are limited. Available studies suggest that perinatal HIV exposure may be associated with alterations in brain development, including structural (30,31), functional (32), and neurometabolite differences between HEU and HU children (33). Further, these findings may contribute to explaining neurodevelopmental differences, as cortical thickness was found to partially mediate the association between maternal HIV and poor language outcomes in children who are HEU from the DCHS at age 2–3 years (34). Magnetic resonance spectroscopy (MRS) provides sensitive, non-invasive measures of brain neurochemistry. In earlier work within the DCHS, maternal and child immune markers were associated with altered neurometabolite ratios in children who are HEU at 2–3 years of age (29), suggesting that immune dysregulation during pregnancy may affect neurometabolite levels in the developing brain. Further research is needed to understand whether these associations persist by school entry, and whether they relate to long-term neurodevelopmental outcomes.

Learning outcomes at school entry are critical indicators of future educational attainment and socioeconomic well-being (35). In South Africa, which has the highest number of children with perinatal HIV exposure globally (36), the Thrive by Five Index Survey showed that fewer than half of preschool-aged children were developmentally on track by 2021 (37). Socioeconomic inequality, malnutrition, and psychosocial adversity are major drivers of these disparities, but less is unknown about the potential contribution of biological influences including maternal HIV and early immune dysregulation (38). Understanding how early-life exposures affect neuroimmune development and shape school readiness in a setting with a high prevalence of perinatal HIV exposure is therefore of both scientific and policy relevance.

Therefore, the aim of this study was to explore neurobiological pathways through which maternal HIV and immune dysregulation may affect child neurodevelopment and school readiness. Using data from a South African birth cohort with high HIV prevalence and socioeconomic adversity, we examined maternal and child immune markers from pregnancy to five years of age, and their associations with brain neurometabolite levels at 6–7 years. We further investigated whether serum immune markers and neurometabolite levels were associated with learning outcomes at school entry, measured using the culturally validated Early Learning Outcomes Measure tool (39).

## Methods Participants

This study was conducted on a sub-group of mother–child dyads enrolled in the DCHS, a population-based birth cohort from the Western Cape, South Africa (40–42). The DCHS investigates early-life determinants of child health and development. The cohort is characterized by a low socioeconomic status and a high prevalence of HIV and associated risk factors. Women were recruited between 2012 and 2015 during their second trimester of pregnancy (gestational age 20–28 weeks) and have been prospectively followed with their children from birth. Eligible participants were aged 18 years or older, planned to attend antenatal care at one of two local public health clinics, and intended to remain in the area. Written informed consent was obtained at enrolment and reaffirmed annually throughout follow-up.

For this study, a subset of children (n=268; 94 HEU, 174 HU) underwent MRI at the Cape Universities Body Imaging Centre (CUBIC). Participants included in this study were children aged 6–7 years who had also undergone neuroimaging at 6–10 weeks and/or 2–3 years of age (42).

Children with medical conditions that could substantially affect neurodevelopment (e.g., congenital anomalies, genetic disorders, neurological disease), low Apgar scores (<7 at 5 minutes), neonatal intensive care admission, maternal substance use during pregnancy, confirmed child HIV infection, or MRI contraindications such as implanted medical devices, were excluded (43). Data on peripheral blood serum marker levels from this neuroimaging subset were available and included maternal samples collected during pregnancy, and child samples collected at ages 6 weeks, 2, 3, and 5 years. In addition to neuroimaging and serum marker data, neurodevelopmental assessments for school readiness at 6–7 years of age were also available.

## Sociodemographic data collection

Maternal HIV status was determined through routine antenatal screening and retesting every 12 weeks through pregnancy, as per Western Cape HIV guidelines at the time (44,45). HIV testing for children who were HIV-exposed followed national early infant diagnosis protocols and included polymerase chain reaction (PCR), rapid antibody, or ELISA testing at 6 weeks, 9 months, and 18 months. Children were confirmed HIV-negative at 18 months or following the cessation of breastfeeding, if this extended beyond that age. HU children were born to mothers with no HIV. All mothers living with HIV received antiretroviral therapy (ART) according to recommendations for prevention of vertical HIV transmission at the time, and HEU infants were given antiretroviral prophylaxis (46). Maternal CD4 counts and viral load during pregnancy were extracted from clinical records and the National Health Laboratory Service database, using the lowest CD4 count and highest viral load recorded within one year before birth and three months postpartum.

Sociodemographic and maternal psychosocial data were collected through structured interviews between 28- and 32-weeks’ gestation, using tools adapted from the South African Stress and Health study (40,41). Infant birthweight and growth indicators were calculated using WHO Z-scores (47). Maternal alcohol consumption during pregnancy was evaluated with the Alcohol, Smoking, and Substance Involvement Screening Test (42). Smoking status during pregnancy was self-reported and assessed using the same tool. Depressive symptoms were measured with the Edinburgh Postnatal Depression Scale (41). All variables were converted into binary categories using established thresholds.

## Immune assays

Maternal serum samples were collected during the second trimester of pregnancy (26–28 weeks’ gestation), while child samples were obtained at 6 weeks, 2, 3, and 5 years of age (40). A cytokine panel consisting of granulocyte-macrophage colony-stimulating factor (GM-CSF), interferon gamma (IFN-γ), interleukins IL-1β, IL-2, IL-4, IL-5, IL-6, IL-7, IL-8, IL-10, IL-12p70, IL-13, and tumor necrosis factor alpha (TNFα) was quantified using a commercially available Milliplex® Luminex 13-plex assay (#HSTCMAG28SPMX13; Merck), following the manufacturer’s protocol (19). Samples were run on a Bio-Plex 200 Luminex platform (Bio-Rad). Additional markers were analysed via enzyme-linked immunosorbent assay (ELISA; R&D Systems, Minneapolis, USA), including soluble monocyte activation markers (sCD14 and sCD163), neutrophil gelatinase-associated lipocalin (NGAL), matrix metalloproteinase-9 (MMP-9), and chitinase-3-like protein 1 (YKL-40). All assays were performed blinded and in duplicate (19).

## Magnetic Resonance Spectroscopy protocol

Children enrolled in the neuroimaging sub-study underwent a multimodal MRI protocol without sedation at the Cape Universities Body Imaging Centre (CUBIC), between August 2018 and July 2022. Neuroimaging was performed using a Siemens 3T Skyra MRI scanner (VE11) equipped with a 32-channel head coil (43). Upon arrival at CUBIC, families were welcomed into a child-friendly environment. Study staff provided detailed explanations of the MRI procedures, obtained written informed consent and child assent, conducted an MRI safety screening, and played an educational video to familiarize families with the scanning process. Before entering the scanner, children selected a movie to watch during the scan to help them remain relaxed and still.

High-resolution structural images were acquired using a T1-weighted MPRAGE sequence (field of view 256×256mm²; TR=2500ms; TI=1000ms; TE=3.35ms; bandwidth 240LHz/Px; voxel resolution 1×1×1mm³). Single-voxel ¹H-MRS acquisitions (25×25×25mm³) were positioned over midline parietal grey matter and left parietal white matter. Spectroscopy data were obtained using a point-resolved spectroscopy sequence (TE=30ms; TR=2000ms; 128 averages; bandwidth 1200Hz; vector size 1024), with chemical shift selective pulses for water suppression. A non-water-suppressed reference scan was also acquired for absolute quantification. Automatic voxel shimming was applied, with manual shimming adjustments made as needed to optimize spectral resolution and minimize linewidth.

## Magnetic Resonance Spectroscopy data processing

Spectral data were analysed using LCModel software (version 6.2), which provided frequency alignment, eddy current and baseline corrections, and quantified both ratios to total creatine and absolute concentrations (48). A standardized *in vitro* basis set was used to fit neurometabolite peaks and derive estimates for glutamate, myo-inositol, total choline, and n-acetyl-aspartate. As the primary excitatory neurotransmitter, glutamate was interpreted as a neurometabolite of excitatory neuronal function; myo-inositol as a proxy for glial activation and neuroinflammation; total choline as reflecting membrane turnover and white matter maturation/myelination; and N-acetyl-aspartate as a surrogate marker of neuronal/axonal integrity (49–51). Tissue segmentation was performed using Statistical Parametric Mapping (SPM12) to estimate voxel composition (grey matter, white matter, cerebrospinal fluid) and to adjust for partial volume effects and water content. Poor quality spectra were excluded (e.g., those showing lipid contamination, abnormal baselines, or visual artifacts). Additional exclusion criteria were full width at half maximum above 0.09Lppm, a signal-to-noise ratio below 15, and Cramér-Rao Lower Bounds greater than 20%. Spectra that had been acquired with a TE longer than 30ms or lacked a water reference were also excluded.

## Early Learning Outcome Measures

School readiness was assessed using the Early Learning Outcome Measure (ELOM) assessment, an age-validated, population-level standardized tool developed for South African children aged 50–69 months (39). The tool evaluates five key domains aligned with the South African early childhood curriculum: gross motor skills, fine motor and visual-motor integration, emergent mathematics and numeracy, cognitive and executive functioning, and emergent language and literacy. Trained research staff administered the ELOM assessment to children in their mother tongue (isiXhosa or Afrikaans) at local community centres near study clinics. ELOM scores were used to capture individual-level developmental profiles and evaluate early learning outcomes in relation to HIV exposure status and neuroimmune findings.

## Statistical analysis

Descriptive statistics for maternal and child sociodemographic data were summarized as medians with interquartile ranges (IQR) for continuous variables, and as frequencies and percentages for categorical variables. The distribution of continuous data was assessed visually using histograms.

To compare characteristics between HEU and HU children, independent *t*-tests or Wilcoxon rank-sum tests were performed, according to data distribution. Categorical comparisons were conducted using Chi-square tests.

Serum marker concentrations were log-transformed prior to analysis. Neurometabolite levels were scaled to approximate normality, and group differences were assessed using independent *t*-tests. Cross-sectional comparisons between groups were conducted using independent *t*-tests or Wilcoxon rank-sum tests, depending on the distribution of each variable. Longitudinal trajectories of child peripheral blood markers from 6 weeks to 5 years of age were analysed using linear mixed-effects models (LMMs) with maximum likelihood estimation. Differences in ELOM scores between groups were also evaluated using independent *t*-tests.

To investigate whether serum marker trajectories predicted neurometabolite levels at 6–7 years, child-specific trajectory features were derived from the LMMs. Best linear unbiased predictions (BLUPs) were extracted for each child and combined with fixed effects to obtain child-specific coefficients. These coefficients described linear slopes, as well as higher-order terms capturing potential non-linearities. Slopes were estimated for children with at least one valid serum marker measurement and were then used as predictor variables in linear models.

Linear regression models with robust standard errors were used to examine three sets of cross-sectional associations: maternal/child serum markers and neurometabolite levels at age 6–7 years; neurometabolite levels and ELOM scores; and maternal/child serum markers and ELOM scores. Maternal HIV status was included as an interaction term in all models, and where this interaction was statistically significant, we subsequently fit stratified models by HIV exposure group to generate the estimates presented in tables. All models were corrected for multiple comparisons using the Benjamini–Hochberg (BH) false discovery rate procedure. Associations that remained significant after BH correction were further examined using adjusted linear models. Covariates were selected a priori based on their reported association with neurometabolite or neurodevelopmental outcomes in children, including child age (49–51), sex (52,53), and voxel tissue composition (33).

Mediation analyses were conducted to explore whether serum marker and neurometabolite levels that showed significantly different adjusted associations in HEU and HU groups mediated the relationship between maternal HIV and child ELOM scores.

Statistical significance was defined as p<0.05 (two-tailed). All analyses were performed using R (version 4.5.0) with RStudio. The full analysis protocol was pre-registered on Open Science Framework (OSF) (54).

## Results

### Cohort and demographic characteristics

Overall, 268 children (94 HEU, 174 HU) from the DCHS underwent MRS at 6–7 years of age. Maternal sociodemographic characteristics were broadly comparable by HIV status, with no group differences in education, monthly household income, employment, relationship status, or anaemia during pregnancy (p>0.05). However, women living with HIV were older at delivery, reported lower antenatal depression rates and lower alcohol and tobacco use during pregnancy, and breastfed for a shorter duration than women without HIV. Birthweight and immunisation coverage were similar between HEU and HU children (**Table 1**).

**Table 1.**
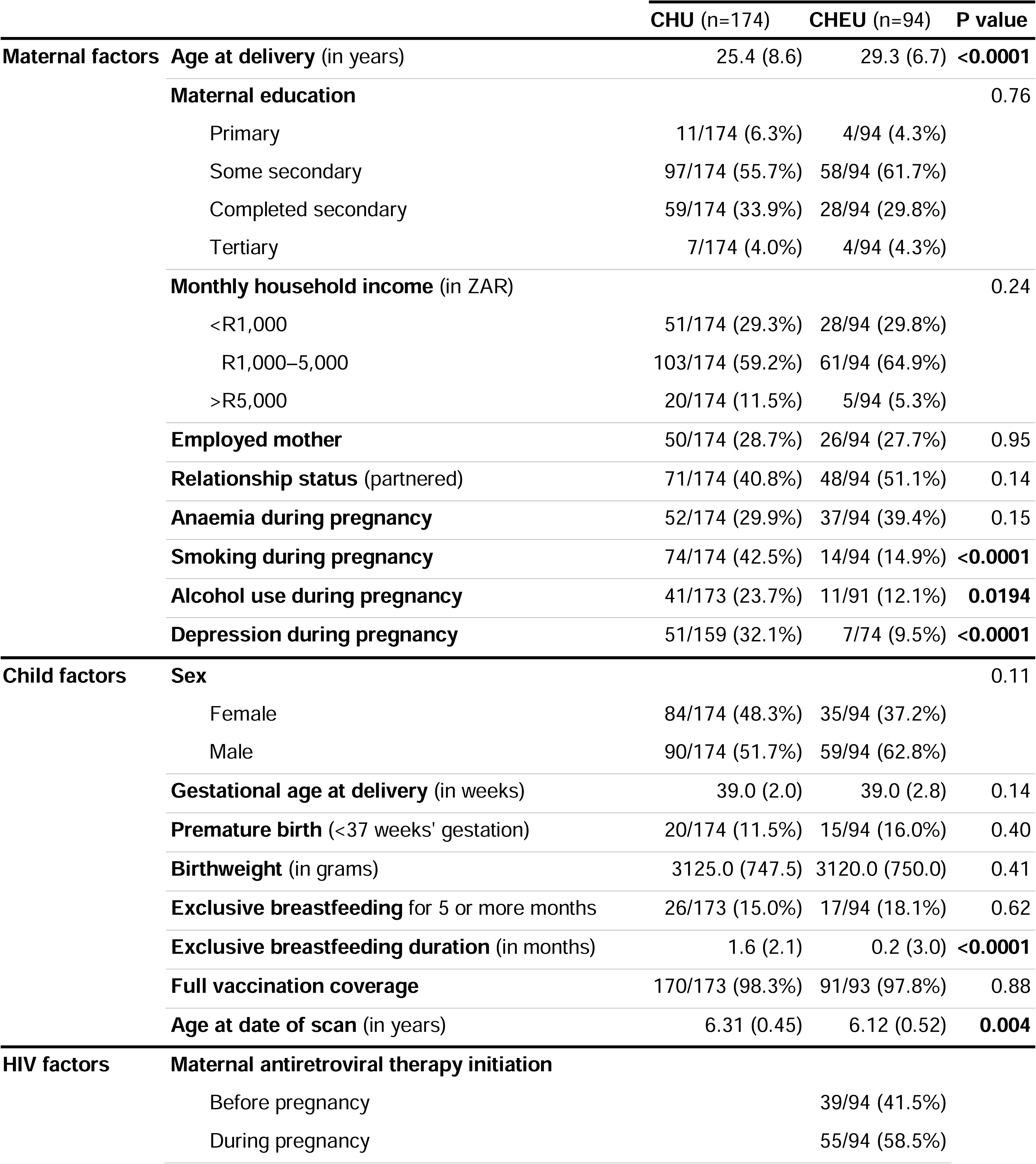
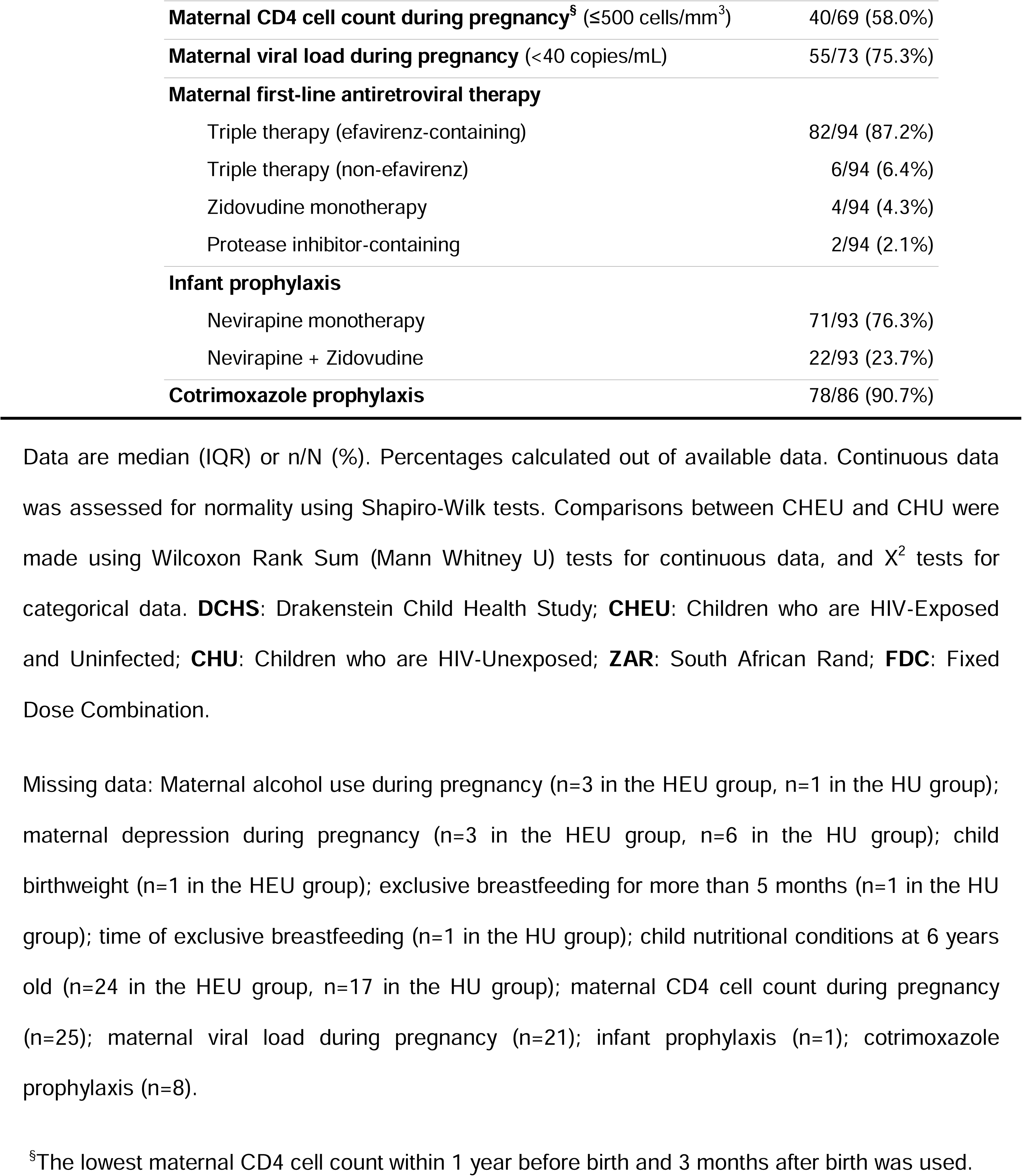
Sociodemographic characteristics of children invited for MRS at age 6–7 years, according to HIV exposure

All mothers living with HIV received ART, initiated either before (41.5%) or during pregnancy (58.5%); most (87.2%) received an efavirenz-based triple regimen. All HEU infants received nevirapine prophylaxis (76.3% monotherapy; 23.7% with zidovudine), and 90.7% received cotrimoxazole.

### Maternal serum immune marker concentrations during pregnancy

Peripheral blood serum immune markers were measured at 26–28 weeks’ gestation in 216 mothers (80.6%, 83 living with HIV, 133 without HIV) of the 268 whose children were invited for neuroimaging. After BH correction, mothers living with HIV had higher levels of monocyte activation marker sCD14 (p=0.002), and lower pro-inflammatory cytokine GM-CSF (p=0.007) and neuroinflammatory NGAL (p=0.018) and MMP-9 (p=0.0006) than those without HIV. Lower IL-13 was observed in mothers with HIV before BH correction only (uncorrected p=0.0196) (**Supplementary Figure 1**, **Supplementary Table 1**).

### Child serum immune marker trajectories

Longitudinal trajectories from 6 weeks to 5 years were modelled in 218/268 children (81.3%, 85 HEU, 133 HU) from the neuroimaging sub-study with one or more serum measurements.

In children who are HU, most markers showed age-related changes across early childhood (**Supplementary Table 2**). Perinatal HIV exposure was associated with distinct developmental profiles for several markers (**Figure 1**). Compared to HU peers, children who are HEU had higher baseline sCD14 (β=0.09, p=0.026) with a divergent non-linear trajectory over time (β=−0.13, p=0.026). IL-5 showed a declining pattern in children who are HEU (β=−0.44, p=0.003) rather than the increase seen in HU peers (β=0.60, p=0.003). IL-1β was lower at baseline in children who are HEU (β=−0.16, p=0.032) but increased linearly (β=0.27, p=0.015), compared to the non-linear trajectory in the HU group. GM-CSF (β=−0.26, p=0.006) and IL-10 (β=−0.18, p=0.020) were lower at baseline, and showed no significant temporal changes, in children who are HEU. YKL-40 also showed an HIV exposure-specific pattern, with a non-linear increase in children who are HEU (β=0.18, p=0.041) contrasting with a decline in HU peers (β=−0.23, p=0.003). Cross-sectional group differences at each timepoint are reported in **Supplementary Table 1**.

**Figure 1.**
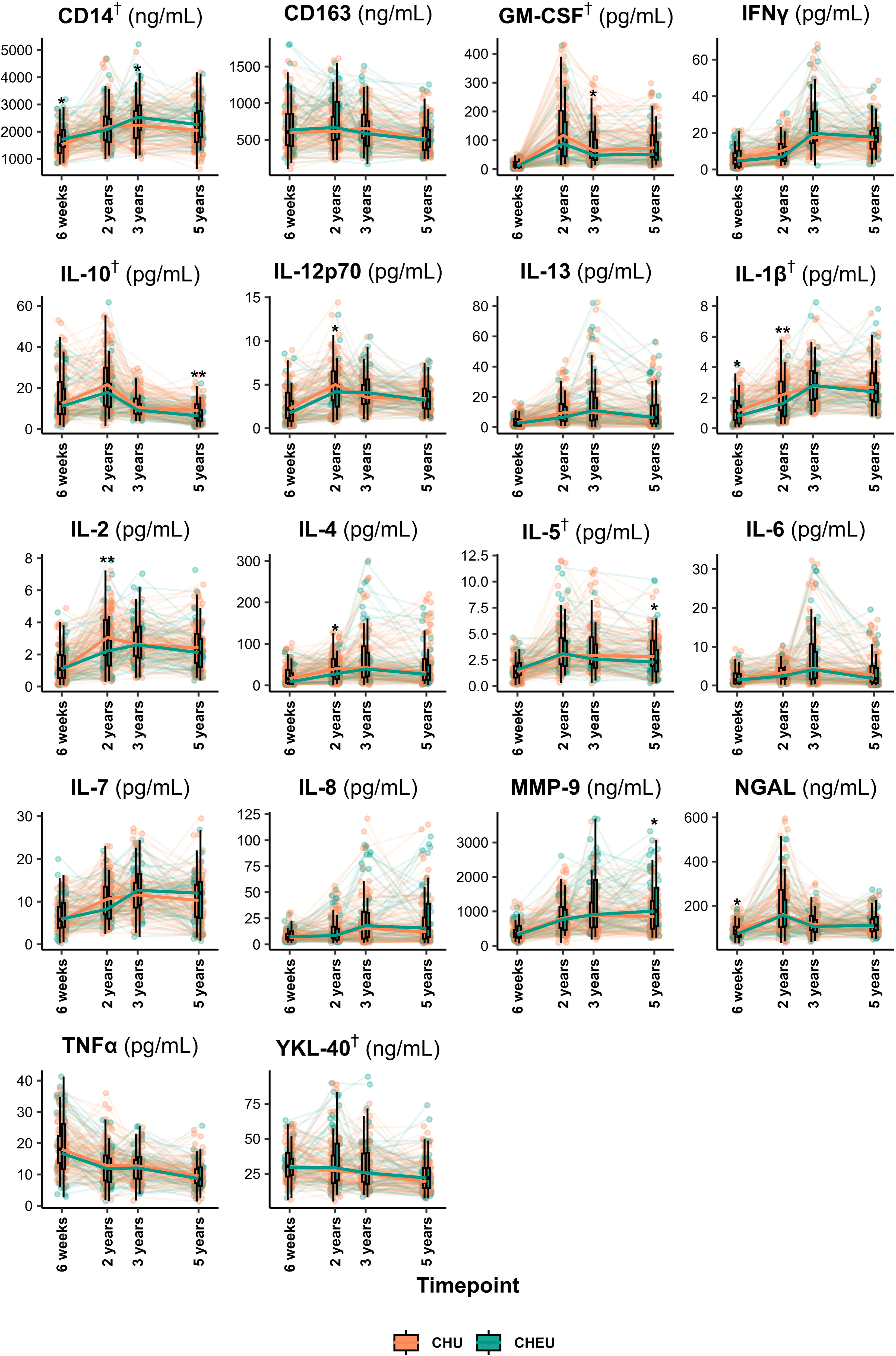
Longitudinal trajectories of child peripheral blood serum biomarkers from 6 weeks to 5 years of age. Trajectories of 18 immune and inflammatory markers (pg/mL or ng/mL), stratified by maternal HIV status. Each panel displays individual child-level measurements, connected across timepoints where available, and group medians with interquartile ranges (boxplots). Group medians across timepoints are joined by thick lines. Data from children with available marker measurements at one or more timepoints were included. Asterisks (*) indicate significant group differences at specific timepoints (*t*-tests or Wilcoxon tests), and daggers (†) denote significant group differences in overall longitudinal trajectories (linear mixed-effects models). **CHEU**: Children who are HIV-Exposed and Uninfected (teal); **CHU**: Children who are HIV-Unexposed (orange).

### Child neurometabolite levels at 6**–**7 years

After excluding low-quality spectra, usable creatine-referenced neurometabolite ratio data were available for 188 children in midline parietal grey matter (57 HEU, 131 HU) and 163 children in left parietal white matter (47 HEU, 116 HU). Absolute neurometabolite estimates were available for 184 children in grey matter (57 HEU, 127 HU) and for 90 children in white matter (16 HEU, 74 HU). The smaller sample for left parietal white matter absolute concentrations reflects exclusion of scans with unreliable water-referenced quantification following an acquisition inconsistency.

Children who are HEU had lower glutamate ratios to creatine in the midline parietal grey matter (p=0.0308) and lower total choline ratios in the left parietal white matter (p=0.0295), than HU peers (**Figure 2**, **Supplementary Table 3**). No other group differences reached statistical significance, although myo-inositol absolute concentrations in left parietal white matter showed a trend toward higher values in children who are HEU (p=0.0646) compared to HU peers (**Supplementary Figure 1**, **Supplementary Table 3**).

**Figure 2.**
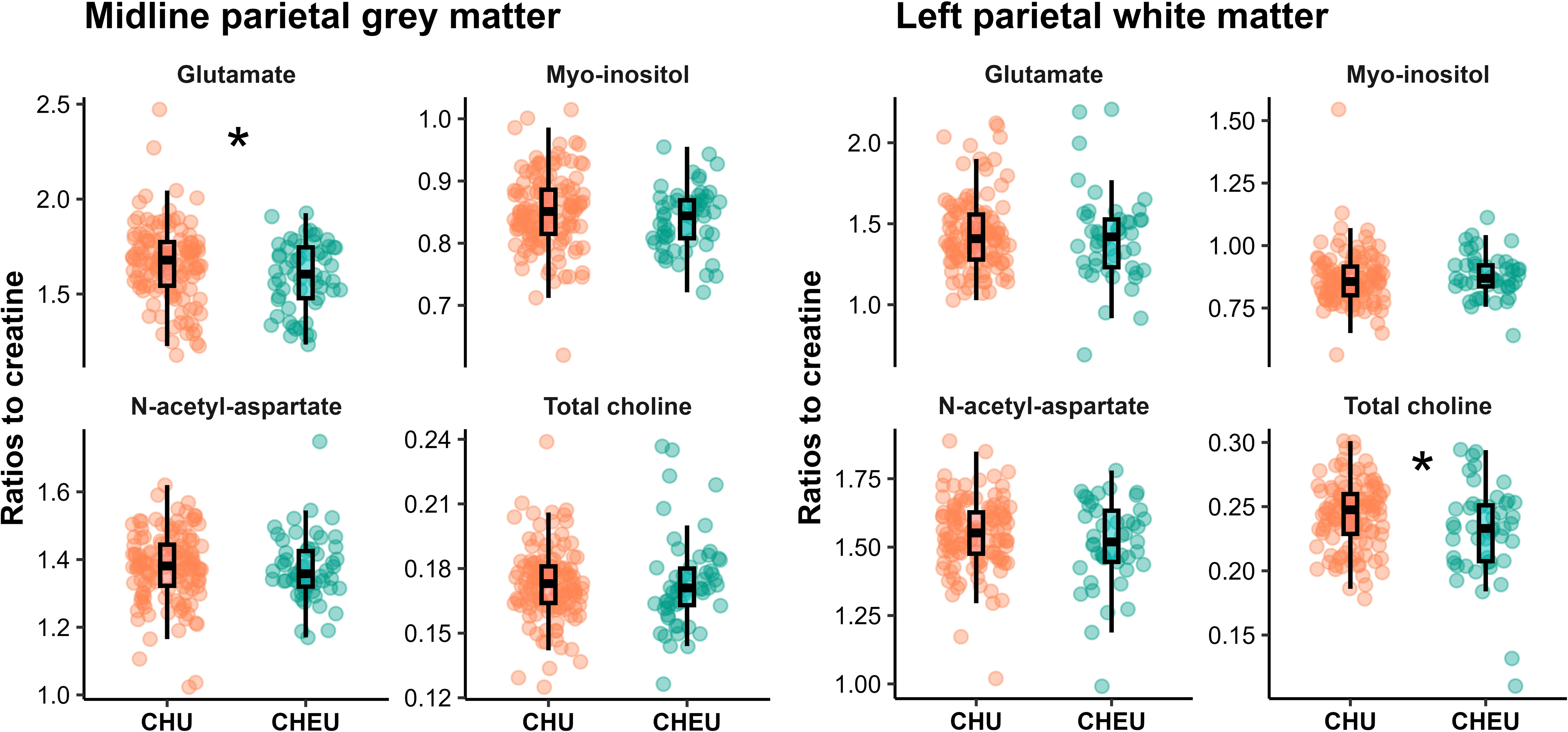
Child neurometabolite ratios to creatine in the parietal brain of children with and without perinatal HIV exposure at age 6–7 years. Raw data plotted. Statistical tests were conducted on scaled data.

### Associations between maternal**/**child serum marker concentrations at different timepoints and child neurometabolite ratios to creatine at 6**–**7 years

**Table 2** summarises associations between serum markers and creatine-referenced neurometabolite ratios that survived BH correction and remained significant in adjusted models; an expanded description is provided in **Supplementary Table 4**. Corresponding results for neurometabolite absolute concentrations are shown in **Supplementary Table 5**.

**Table 2.**
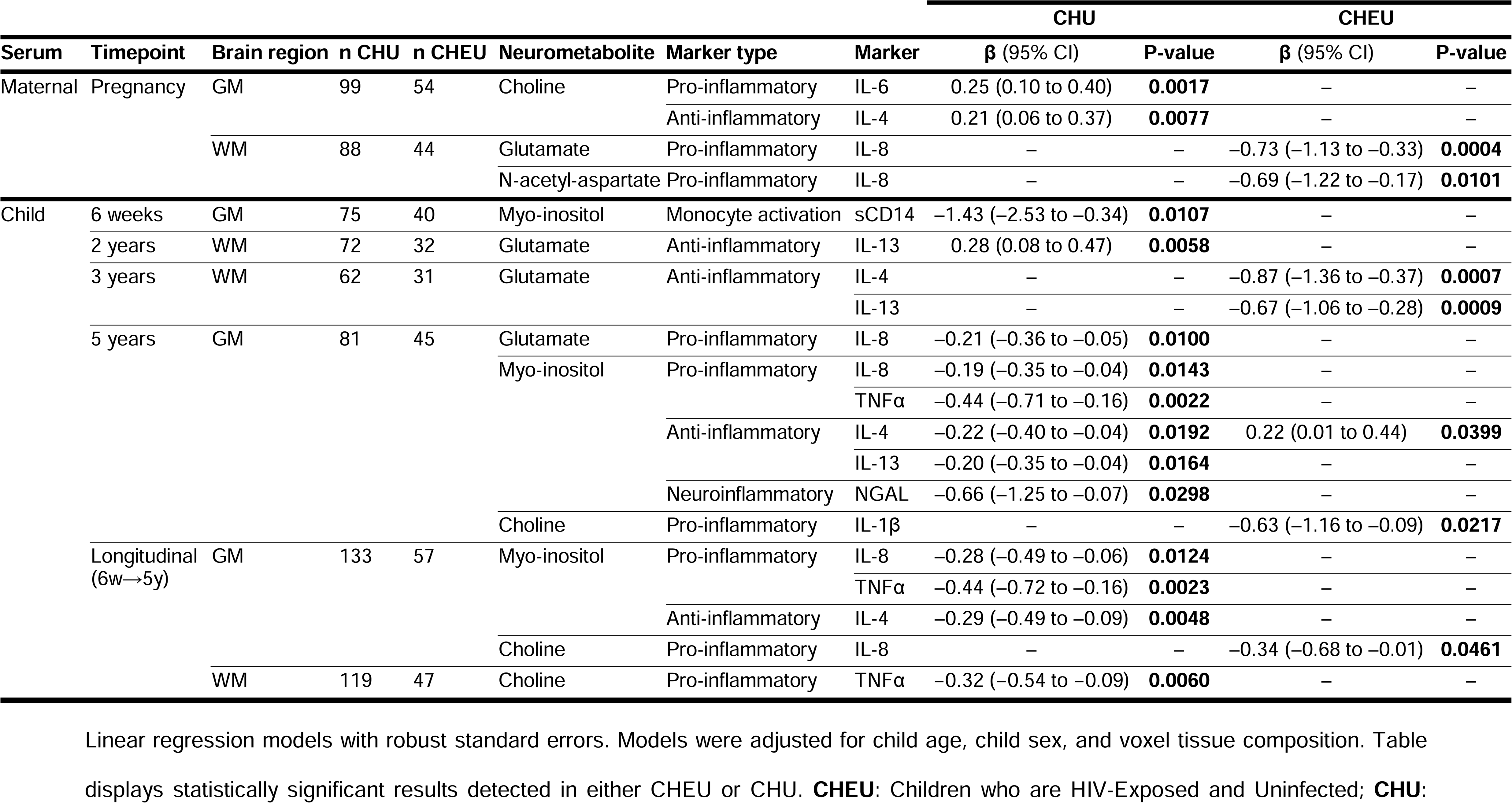
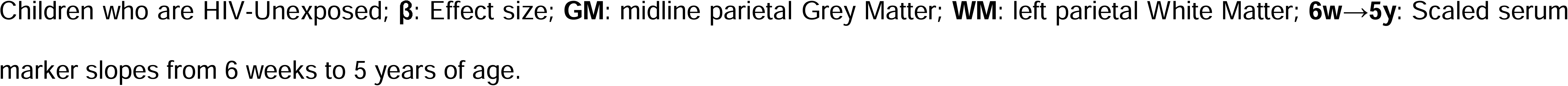
Adjusted linear models exploring associations between maternal/child serum marker levels at different timepoints and child neurometabolite ratios at 6**–**7 years

### Maternal serum marker levels during pregnancy

Maternal cytokines at 26–28 weeks’ gestation showed HIV exposure-specific associations with child neurometabolite ratios: maternal IL-8 was negatively associated with left parietal white matter glutamate ratios (p=0.0004) and N-acetyl-aspartate ratios (p=0.0101) in children who are HEU; maternal IL-6 (p=0.0017) and IL-4 (p=0.0077) were positively associated with midline parietal grey matter total choline ratios in HU peers (**Table 2**).

### Child serum markers

In HU children, trajectories of IL-8 (p=0.0124), TNFα (p=0.0023), and IL-4 (p=0.0048) from 6 weeks to 5 years of age were negatively associated with midline parietal grey matter myo-inositol ratios, and TNFα trajectories were negatively associated with left parietal white matter total choline ratios (p=0.0060). In children who are HEU, IL-8 trajectories were negatively associated with midline parietal grey matter total choline ratios (p=0.0461) (**Table 2**).

In cross-sectional analyses, sCD14 was negatively associated with parietal grey matter myo-inositol ratios in HU children (p=0.0107), and IL-13 at age 2 years was positively associated with left parietal white matter glutamate ratios (p=0.0058). In children who are HEU, IL-4 (p=0.0007) and IL-13 (p=0.0009) at age 3 years were negatively associated with left parietal white matter glutamate ratios. At age 5 years, multiple inflammatory markers were negatively associated with midline parietal grey matter myo-inositol ratios in HU children, including IL-8 (p=0.0143), TNFα (p=0.0022), IL-4 (p=0.0192), IL-13 (p=0.0164), and NGAL (p=0.0298). Also in the midline parietal grey matter voxel, age-5 IL-4 in children who are HEU was positively associated with myo-inositol ratios (p=0.0399), and IL-1β was negatively associated with total choline ratios (p=0.0217) (**Table 2**).

### Sensitivity analyses

To account for group differences in antenatal exposures, sensitivity analyses additionally adjusted for sociodemographic and psychosocial variables that differed between HEU and HU groups (**Table 1**). Adjustment for maternal alcohol use during pregnancy attenuated the association between age-5 IL-4 and grey matter myo-inositol ratios in children who are HEU; all other associations were unchanged (**Supplementary Table 6**).

### Associations between maternal**/**child serum marker concentrations at different timepoints and child neurometabolite absolute concentrations at 6**–**7 years

Adjusted models identified several HIV exposure-specific associations between child serum markers and midline parietal grey matter absolute neurometabolite concentrations (in mM) at 6–7 years (**Supplementary Table 5**). In children who are HEU, IL-4 at 6 weeks was negatively associated with total choline levels (β=−0.27, p=0.0310). At age 2 years, sCD14 was negatively associated with glutamate (β=−0.86, p=0.0135) and myo-inositol concentrations (β=−0.74, p=0.0333) in HU children; TNFα was negatively associated with myo-inositol concentrations in children who are HEU (β=−0.47, p=0.0422). At age 3 years, IFN-γ (β=−0.38, p=0.0288) and sCD14 (β=−0.71, p=0.0495) were negatively associated with glutamate concentrations, while IL-5 was positively associated with N-acetyl-aspartate concentrations, in HU children (β=0.28, p=0.0321).

Serum marker association analyses with absolute neurometabolite concentrations in the left parietal white matter are not presented because the small sample size for HEU children (n=16) precluded adequately adjusted models and yielded unstable estimates. As prespecified, only BH-corrected, adjusted associations are reported.

### ELOM scores

Neurodevelopmental data at 6–7 years were available for 172/268 children (64.2%, 56 HEU, 116 HU) from the neuroimaging sub-study. Children who are HEU had lower ELOM language and literacy scores (median 11.2, IQR 5.3) than their HU peers (median 13.6, IQR 6.2) (β=1.95, 95% CI 0.53 to 3.37, p=0.008). No group differences were seen for cognitive development, motor coordination, gross motor development, or emergent mathematics. Overall school readiness scores were lower in children who are HEU (median 65.7, IQR 17.2) compared to HU (median 68.1, IQR 15.5) but did not reach statistical significance (β=1.94, 95% CI −1.92 to 5.93, p=0.33) (**Supplementary Table 7**).

### Associations with maternal serum marker levels during pregnancy

Associations that were significant in unadjusted models, survived BH correction, and remained significant after adjustment for confounders are presented in **Table 3**. In children who are HEU, maternal IL-8 at 26–28 weeks’ gestation was negatively associated with gross motor development at school entry, and sCD163 was positively associated with fine motor coordination. In HU children, maternal IL-2, IL-7, and IL-12p70 in mothers without HIV were negatively associated with cognitive development. In the same group, IFNγ and IL-7 were negatively associated with numeracy and overall school readiness scores.

**Table 3.**
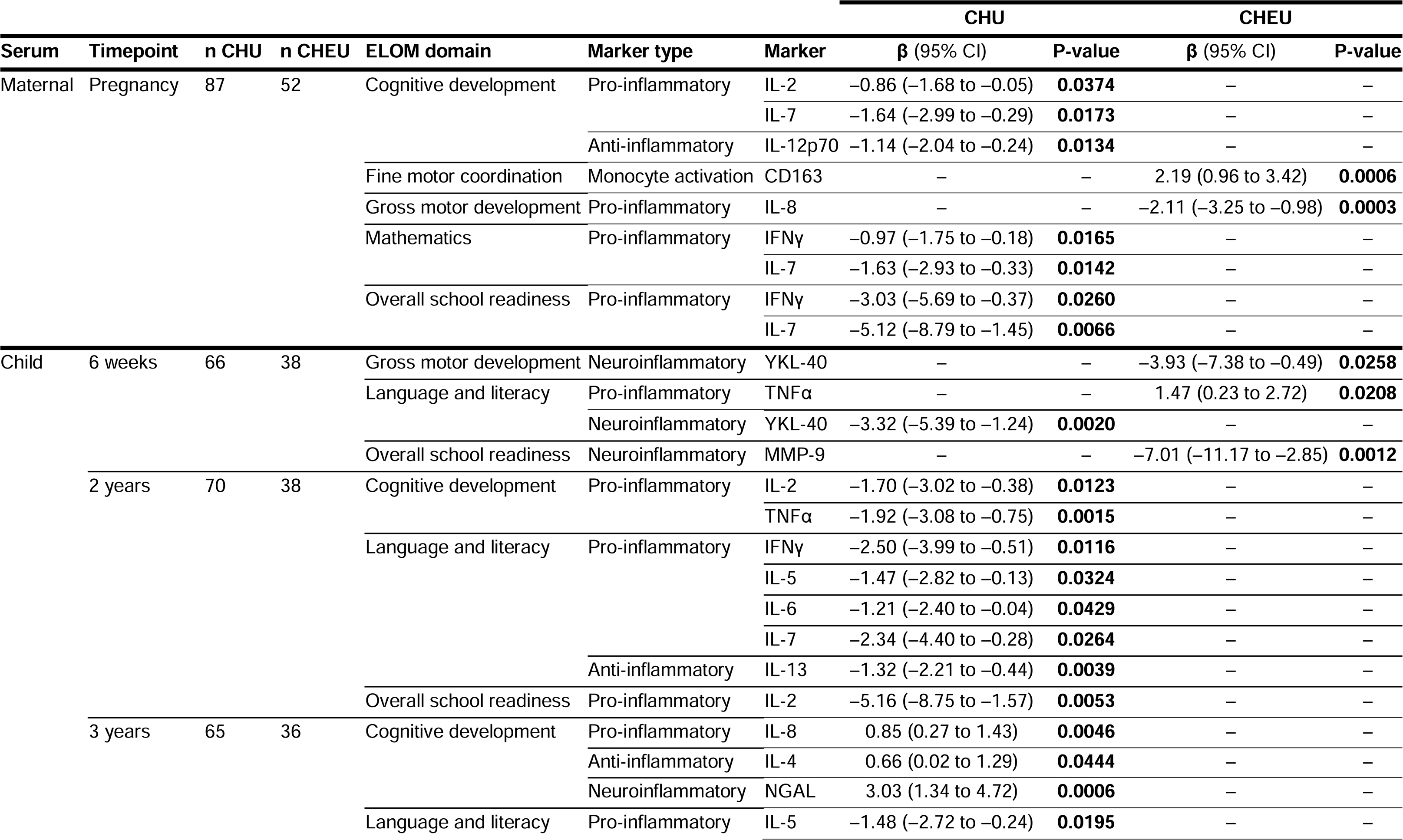
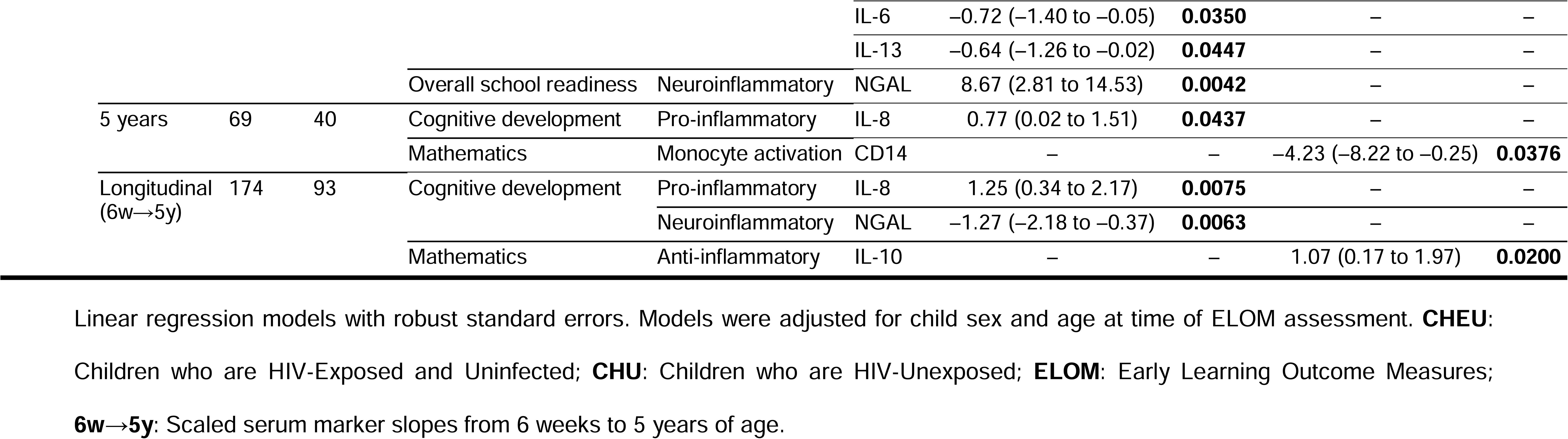
Adjusted linear models exploring associations between maternal and child serum marker levels and early learning outcome measure scores

### Associations with child serum markers

Longitudinally, IL-10 trajectories in children who are HEU were positively associated with numeracy scores. In HU children, cognitive scores were positively associated with IL-8 trajectories and negatively associated with NGAL trajectories (**Table 3**).

In cross-sectional analyses, YKL-40 at 6 weeks in children who are HEU was negatively associated with gross motor development, TNFα positively associated with language and literacy, and MMP-9 negatively associated with overall school readiness scores. In HU children, YKL-40 was negatively associated with language and literacy scores. At 2 years, IL-2 and TNFα in HU children were negatively associated with cognitive scores; IFNγ, IL-5, IL-6, IL-7, and IL-13 were negatively associated with language and literacy; and IL-2 was negatively associated with overall school readiness scores. At 3 years, IL-8, IL-4, and NGAL in HU children were positively associated with cognitive scores, and NGAL was positively associated with overall school readiness scores; conversely, IL-5, IL-6, and IL-13 were negatively associated with language and literacy scores. At 5 years, sCD14 in children who are HEU was negatively associated with numeracy scores, while in HU children, IL-8 was positively associated with cognitive scores.

### Associations with child neurometabolite levels at 6–7 years

In HU children, left parietal white matter total choline ratios to creatine were associated with higher ELOM language and literacy scores (β=0.08, 95% CI 0.04 to 0.13, p=0.0003) and higher overall school readiness scores (β=0.03, 95% CI 0.02 to 0.05, p<0.0001). This was not observed in children who are HEU. No other associations remained significant after BH correction and adjustment.

In unadjusted analyses, left parietal white matter total choline absolute concentrations were associated with ELOM language and literacy scores (β=0.08, 95% CI 0.02 to 0.15, p=0.007) and overall school readiness (β=0.03, 95% CI 0.01 to 0.05, p=0.003) in HU children.

### Mediation effects

Mediation analyses were performed to assess whether serum marker and neurometabolite levels that showed significant associations in either HEU or HU groups mediated the relationship between maternal HIV and child ELOM scores (see **Mediation Analysis** in **Supplementary Files**). Separate models were tested for each marker–neurometabolite–score pathway, all of which demonstrated adequate fit based on model indices. None of the pathways tested demonstrated significant mediation in this study.

## Discussion

In this longitudinal study, we build on prior DCHS findings in early childhood (29) to demonstrate that neuroimmune associations remain altered in children who are HEU at school age. Mothers living with HIV exhibited an antenatal immune profile characterized by higher sCD14 and lower MMP-9, NGAL, and GM-CSF levels, while children who are HEU showed altered trajectories of monocyte activation markers (sCD14), cytokines (GM-CSF, IL-1β, IL-5, IL-10), and neuroinflammatory markers (YKL-40) from 6 weeks to 5 years of age compared to their HU peers. Neurometabolite differences were detected at school entry, with children who are HEU showing lower glutamate ratios to creatine in the midline parietal grey matter and lower total choline ratios in the left parietal white matter compared to their HU peers, alongside trends toward higher white matter myo-inositol absolute concentrations. Multiple HIV exposure-specific associations were observed between maternal/child serum markers and neurometabolite ratios to creatine at 6–7 years. Notably, the pattern shifted from maternal marker associations in early life (29) to predominantly child marker associations by school entry. Overall, children who are HEU performed worse on language assessments while, in HU peers, higher left parietal white matter choline ratios to creatine associated with better language scores, suggesting a relationship between neuroimmune alterations and school readiness.

The antenatal immune differences observed between mothers living with and without HIV, including higher sCD14 levels, support previous reports in pregnant women with HIV (21,55) as well as in children with perinatal HIV infection (56) or exposure (24,55). This highlights the potential importance of monocyte activation pathways in peripheral inflammation in both mothers living with HIV and their children who are HEU. Lower levels of the neuroinflammatory marker MMP-9, as observed in mothers with HIV in our cohort, have also been reported in adults with HIV (57), while neuroinflammatory NGAL has been linked to cognitive impairment in adult patients living with HIV (57,58). YKL-40, which displayed different trajectories in HEU and HU children in our study, has been associated with language and gross motor delays in Ugandan children who are HEU at 18 months of age (59).

Neurometabolite differences at 6–7 years in our cohort, including lower glutamate ratios to creatine in the midline parietal grey matter of HEU children, align with previous reports of decreased grey matter glutamate concentrations in South African children who are HEU at older ages (60,61), and may reflect disruptions in synaptic efficacy (49–51) or impaired astrocyte–neuron glutamate cycling (62). Similarly, lower total choline ratios to creatine observed in the left parietal white matter of HEU children in our cohort are consistent with previously described white matter microstructural alterations in perinatal HIV exposure (32,63,64) and may represent delayed myelination or reduced membrane turnover (49–51). White matter development begins in the fetal period, maturation accelerates during early-to-mid childhood, and continues into early adulthood before reaching mature peak volume (65). White matter remains highly plastic and vulnerable during these stages; therefore, our findings may reflect early disruptions in myelination and membrane turnover persisting into school age. Importantly, we also observed a trend toward higher white matter myo-inositol concentrations in HEU children, consistent with the neuroinflammatory pattern previously described in this cohort at an earlier age (33).

Neuroimmune associations in our study align with and extend earlier DCHS work at age 2–3 years, but with evidence of a developmental shift: maternal serum markers were more strongly associated with neurometabolites in early life (29), whereas by school entry associations were dominated by child immune markers, as exemplified by IL-8. Maternal IL-8 during pregnancy was positively associated with myo-inositol ratios to creatine in children who are HEU at age 2–3 years (29). In the current study at 6–7 years, child IL-8 trajectories showed differential associations with myo-inositol and choline ratios by maternal HIV status, and IL-8 at 5 years was associated with glutamate and myo-inositol ratios to creatine in children who are HEU. IL-8 is a pro-inflammatory chemokine that recruits neutrophils and modulates inflammation during pregnancy (66), and has been reported to be altered in pregnant women with HIV and their HEU children in Kenya (67). In the brain, IL-8 is produced by microglia and other brain cells in response to inflammatory cues and can modulate neurogenesis, synaptic plasticity, and neuroinflammatory tone during gestation and early infancy (68). Although neuroprotective at physiological levels, excessive or prolonged elevations may become neurotoxic and impair synaptic development (69–72). Consistent with this, higher perinatal IL-8 in neonatal cohorts has been linked to altered white matter connectivity (68,69) and increased risks of neurodevelopmental impairment (70–72). In children who are HEU, the elevated myo-inositol pattern observed at 2–3 years (33) may therefore reflect early glial activation, with downstream effects later manifesting as altered excitatory neurotransmission (glutamate) and membrane turnover (choline) by school entry. These findings highlight IL-8 as a potential marker of persistent neuroimmune alterations in the context of perinatal HIV exposure across developmental stages.

Beyond HIV exposure-specific effects, our findings contribute to the understanding of typical neuroimmune development in high-burden settings. Among HU children, maternal IL-4 and IL-6 positively associated with total choline ratios to creatine in the parietal grey matter, a pattern compatible with typical neurometabolite development. IL-4 is an anti-inflammatory cytokine that supports tissue integrity during fetal brain development (11,73). IL-6 has pleiotropic roles: its classic signalling pathway supports homeostasis, while excessive signalling mediates pro-inflammatory cascades during maternal immune activation (9,10,66,74). Balanced IL-4 and IL-6 signalling during pregnancy may therefore promote optimal myelination and membrane turnover (11), processes for which total choline serves as a surrogate marker (49–51). These divergent neuroimmune relationships in HEU and HU groups are consistent with models of maternal immune activation that propose a disruption of fetal neuroimmune homeostasis in response to prenatal inflammation (7,9).

The developmental trajectories of child immune markers further highlight HIV exposure-specific patterns. In HU children, steeper declines from infancy to age five in pro-inflammatory IL-8 and TNFα, as well as in the anti-inflammatory cytokine IL-4 were associated with lower parietal grey matter myo-inositol ratios to creatine. These results suggest that, in HU children, peripheral cytokines decrease over early childhood alongside reduced glial activation in the brain, consistent with typical neuroimmune homeostasis (11,73). While transient TNFα signalling is essential for normal synaptic pruning, sustained elevation can activate microglia and astrocytes and upregulate pro-inflammatory pathways (8,10,73,74). Accordingly, the natural developmental decline in TNFα during immune maturation (75) is consistent with decreasing myo-inositol as microglial activity subsides and cortical networks stabilize (8,11). Overall, our HU findings align with a model in which the peripheral immune system and brain microglia mature together, supporting healthy brain development (7,11).

In contrast, children who were HEU showed a markedly different pattern, where steeper IL-8 slopes across early childhood predicted lower choline ratios to creatine in the parietal grey matter at 6–7 years, and several single-timepoint associations were inverted relative to HU. Persistent or dysregulated IL-8 signalling in this group may interfere with typical brain development, leading to inefficient or maladaptive membrane turnover (49–51). This is congruent with evidence that IL-8 dysregulation links perinatal systemic inflammation to atypical brain maturation (68,69) and later cognitive risk (70–72), and it supports a model in which perinatal HIV exposure may alter glial sensitivity to peripheral inflammatory pathways. This further supports IL-8 as a potential marker of ongoing neuroimmune alterations in children who are HEU from pregnancy through school age.

School readiness analyses provided a functional real-world perspective on these biological findings. Among HU children, higher left parietal white matter total choline ratios to creatine were associated with higher language and overall school readiness scores, consistent with myelination and membrane turnover processes supporting language networks (49–51,76). These associations were absent in HEU peers, who showed both lower white matter choline ratios to creatine and lower language scores despite broadly similar socioeconomic profiles, echoing wider literature (5). In children who are HEU, serum marker–ELOM associations were mainly for pro-inflammatory cytokines (i.e., IL-8, TNFα) monocyte activation (i.e., sCD14, sCD163) and neuroinflammatory markers (i.e., MMP-9, YKL-40). This suggests that persistent systemic inflammation may be associated with neurodevelopmental outcomes, particularly in domains reliant on developing white matter networks (76). Notably, the lack of significant mediation by any single marker or neurometabolite indicates that maternal HIV likely influences language development through multiple, partly overlapping pathways (including immune programming, ART exposure, perinatal environmental factors, and ongoing psychosocial stressors) rather than a single biological mechanism. In line with this, studies have shown that socioeconomic adversity and prenatal environmental stressors modulate gestational IL-8 concentrations and, in turn, white matter development and early cognitive outcomes in children (69,72).

Key strengths of this study include its prospective, longitudinal design, with data collection from pregnancy through school entry in a well-characterized South African birth cohort. The repeated measurement of child immune markers across multiple stages enabled trajectory modelling. The integration of peripheral immune data with *in vivo* measures of neurometabolites via MRS is another major strength, offering a unique window into neuroimmune associations in the context of perinatal HIV exposure. In addition, culturally validated assessment of school readiness using the ELOM tool allowed us to explore associations between biological findings and functional developmental outcomes in a manner valid for local context. Finally, the study addresses a critical gap by focusing on children who are HEU in a low-resource, high-burden setting, providing evidence that is globally relevant yet grounded in the realities of populations most affected by HIV. Several limitations warrant caution in interpretation. First, although our sample size is larger than many previous neuroimaging studies in children who are HEU, stratification by HIV exposure group and restriction to participants with complete, high-quality MRS and immune data substantially reduced the number of observations contributing to each model. This limited our power to detect modest effects and resulted in wide confidence intervals, particularly for neurometabolite absolute concentrations, which were only available in a small subset. As a consequence, some true associations may have gone undetected. Second, our single-voxel MRS approach focused on parietal regions and therefore cannot capture network-wide heterogeneity or the dynamics of neurotransmitter cycling; future work should incorporate multivoxel spectroscopic imaging alongside complementary neuroimaging modalities to provide a more comprehensive view of brain metabolite spectra. Third, while we adjusted for multiple covariates and conducted sensitivity analyses, residual confounding remains possible given the complex interplay of biological and social factors in this population.

In a setting with high prevalence of perinatal HIV exposure, we show that neuroimmune alterations present in early childhood persist through school age, with HIV exposure-specific associations between peripheral immune markers and neurometabolites, as well as associations with school readiness outcomes. Clinically, this underscores the importance of optimizing antenatal immune health, and the potential use of postnatal immune mediators as prognostic markers for language outcomes as a neurodevelopmental domain of particular vulnerability in children who are HEU (5,6). Mechanistically, integrating cytokine network analyses with MRS indices of astrocytic and microglial function, myelin-sensitive imaging, and longitudinal cognitive phenotyping could clarify pathways linking immune dysregulation to white matter maturation. Future work should also delineate the contributions of specific antiretroviral exposures to these pathways to inform targeted prevention strategies. Replication across other high-burden contexts and deeper mechanistic studies are warranted to validate these findings and understand the trajectories into adolescence.

## Ethics approval and consent

The studies involving human participants were reviewed and approved by the Faculty of Health Sciences, Human Research Ethics Committee, University of Cape Town (401/2009; 525/2012 & 199/2024), by Stellenbosch University (N12/02/0002), and by the Western Cape Provincial Health Research Committee (2011RP45). Written informed consent to participate in this study was provided by the participants’ parent/legal guardian.

## Author Contributions

**CBC**: methodology, formal analysis and interpretation, visualization, writing – original draft, review & editing. **FR**: methodology, formal analysis, supervision, and writing – review & editing. **SW**: methodology, data curation and writing – review & editing. **TK**: methodology, data curation and writing – review & editing. **JA**: methodology, data curation and writing – review & editing. **JR**: methodology, data curation and writing – review & editing. **LB**: methodology, data curation and writing – review & editing. **NH**: project administration and writing – review & editing. **HZ**: conceptualization, methodology, resources, and writing - review & editing. **DS**: conceptualization, methodology, investigation, resources, supervision, and writing – review & editing. **KD**: conceptualization, methodology, investigation, resources, supervision, and writing – review & editing. **PN**: conceptualization, methodology, investigation, data curation, supervision, and writing – review & editing. **CW**: conceptualization, methodology, investigation, data curation, supervision, and writing – review & editing.

All authors approved the final version.

## Data Availability

The Drakenstein Child Health Study is committed to the principle of data sharing. De-identified data will be made available to requesting researchers as appropriate. Requests for collaborations to undertake data analysis are welcome.

More information can be found on our website http://www.paediatrics.uct.ac.za/scah/dclhs. A comprehensive Statistical Analysis Report, as well as the R code used for this study analysis, are openly accessible on OSF (54).

## Acknowledgments

In memory of Professor Dan Stein and Dr Annerine Roos. We show our deep gratitude to the mothers and their children for participating in the Drakenstein Child Health Study. We thank the study staff at Mbekweni, T.C.Newman clinics and Paarl hospital for their hard work and support throughout the duration of the project, and the clinical and administrative staff of the Western Cape Government Health Department at Paarl Hospital and at the clinics for support of the study. We are also grateful for the assistance provided by the Cape Universities Body Imaging Centre (CUBIC) team, particularly Petty Samuels, Mazwi Maishi, and Mariaan Jaftha.

