## Supplementary materials for "Persistent neuroimmune alterations in children who are HIV-exposed but uninfected at age 6–7 years: Associations with language development in a South African birth cohort"

[‡]Shared last authorship

**Keywords**: HIV exposure, cytokine, maternal immune activation, magnetic resonance spectroscopy, brain development, monocyte activation

### Supplementary Tables

### Supplementary Table 1. Cross-sectional group differences in serum marker levels

#### 1.1. Maternal serum markers during pregnancy (2nd–3rd trimester)

| Marker type | Marker | HIV- moms (n=133) | HIV+ moms (n=83) | P value Crude | Corrected |
|---|---|---|---|---|---|
| Pro-inflammatory cytokines (pg/mL) | **GM-CSF** | **49.01 (61.59)** | **32.26 (30.86)** | **0.001** | **0.007** |
| | IFNγ | 9.69 (8.19) | 10.48 (10.86) | 0.68 | 0.77 |
| | IL-1β | 1.72 (1.37) | 1.84 (1.71) | 0.66 | 0.77 |
| | IL-2 | 2.26 (2.47) | 2.15 (2.57) | 0.41 | 0.56 |
| | IL-5 | 2.28 (1.73) | 2.58 (2.55) | 0.65 | 0.78 |
| | IL-6 | 2.48 (2.6) | 1.66 (2.5) | 0.10 | 0.27 |
| | IL-7 | 10.6 (6.91) | 9.7 (8.21) | 0.88 | 0.88 |
| | IL-8 | 3.96 (4.96) | 3.4 (3.76) | 0.31 | 0.56 |
| | TNFα | 5.44 (2.84) | 5.67 (4.28) | 0.12 | 0.31 |
| Anti-inflammatory cytokines (pg/mL) | IL-4 | 38.13 (44.31) | 22.11 (42.86) | 0.05 | 0.16 |
| | IL-10 | 12.02 (15.78) | 10.95 (12.11) | 0.31 | 0.56 |
| | IL-12p70 | 3.73 (3.35) | 3.73 (3.07) | 0.35 | 0.56 |
| | **IL-13** | **6.04 (7.1)** | **3.34 (5.04)** | **0.0196** | 0.07 |
| Monocyte activation markers (ng/mL) | **CD14** | **1582.92 (742.01)** | **1994.69 (802.75)** | **0.0002** | **0.002** |
| | CD163 | 555.36 (388.43) | 545.82 (360.69) | 0.88 | 0.93 |
| Neuroinflammatory markers (ng/mL) | **NGAL** | **169.15 (145.61)** | **138.41 (123.55)** | **0.004** | **0.018** |
| | **MMP-9** | **1226.05 (1022.48)** | **793.61 (910.7)** | **0.00003** | **0.0006** |
| | YKL-40 | 36.41 (30.8) | 34.37 (44.16) | 0.47 | 0.60 |

Group data is presented as raw median (IQR). Statistical tests were conducted on log-scaled data. Analyses were corrected for multiple comparisons using the Benjamini-Hochberg procedure.

**1.2. Infant serum markers** at 6 weeks of age

| Marker type | Marker | CHU (n=100) | CHEU (n=56) | P value Crude | Corrected |
|---|---|---|---|---|---|
| Pro-inflammatory cytokines (pg/mL) | GM-CSF | 14.41 (16.69) | 11.2 (13.63) | 0.09 | 0.34 |
| | IFNγ | 5.74 (5.64) | 4.78 (9.57) | 0.53 | 0.83 |
| | **IL-1β** | **1.15 (1.3)** | **0.85 (1.21)** | **0.0231** | 0.14 |
| | IL-2 | 1.11 (1.43) | 1.1 (1.39) | 0.67 | 0.83 |
| | IL-5 | 1.38 (1.32) | 1.72 (1.31) | 0.09 | 0.34 |
| | IL-6 | 1.58 (3.05) | 1.4 (2.41) | 0.51 | 0.83 |
| | IL-7 | 5.75 (5.07) | 5.92 (5.68) | 0.49 | 0.83 |
| | IL-8 | 6.95 (6.8) | 7.5 (9.38) | 0.99 | 0.99 |
| | TNFα | 18.19 (9.74) | 17.13 (16.73) | 0.35 | 0.70 |
| Anti-inflammatory cytokines (pg/mL) | IL-4 | 16.92 (32.81) | 8.32 (25.12) | 0.20 | 0.52 |
| | IL-10 | 12.46 (15.63) | 11.09 (12.34) | 0.60 | 0.83 |
| | IL-12p70 | 2.24 (3.01) | 1.9 (2.37) | 0.14 | 0.41 |
| | IL-13 | 3.39 (5.34) | 2.95 (4.94) | 0.99 | 0.99 |
| Monocyte activation markers (ng/mL) | **CD14** | **1521.49 (652.66)** | **1716.6 (627.05)** | **0.0109** | 0.20 |
| | CD163 | 603.29 (427.77) | 634.12 (436.03) | 0.67 | 0.86 |
| Neuroinflammatory markers (ng/mL) | **NGAL** | **88.4 (40.28)** | **74.29 (41.35)** | **0.0102** | 0.13 |
| | MMP-9 | 364.84 (252.23) | 342.55 (483.68) | 0.74 | 0.83 |
| | YKL-40 | 30.65 (18.12) | 29.49 (12.14) | 0.47 | 0.73 |

Group data is presented as raw median (IQR). Statistical tests were conducted on log-scaled data. Analyses were corrected for multiple comparisons using the Benjamini-Hochberg procedure.

#### **1.3. Child serum markers** at age 2 years

| **Marker type** | **Marker** | **CHU** (n=107) | **CHEU** (n=60) | **P value** Crude | Corrected |
|---|---|---|---|---|---|
| Pro-inflammatory cytokines (pg/mL) | GM-CSF | 119.97 (147.63) | 91.96 (135.1) | 0.14 | 0.25 |
| | IFNγ | 10.32 (7.11) | 6.98 (8.51) | 0.11 | 0.25 |
| | **IL-1β** | **2.2** (**1.89**) | **1.61** (**1.51**) | **0.0049** | 0.07 |
| | **IL-2** | **3.06** (**2.21**) | **2.21** (**1.59**) | **0.0081** | 0.07 |
| | IL-5 | 3.04 (3.31) | 3.1 (2.43) | 0.48 | 0.57 |
| | IL-6 | 3.42 (2.84) | 2.54 (2.7) | 0.53 | 0.64 |
| | IL-7 | 10.6 (6.5) | 8.28 (5.14) | 0.15 | 0.25 |
| | IL-8 | 9.74 (15.41) | 8.9 (15.04) | 0.29 | 0.38 |
| | TNFα | 13.02 (8.65) | 12.28 (7.64) | 0.29 | 0.42 |
| Anti-inflammatory cytokines (pg/mL) | **IL-4** | **40.47** (**42.65**) | **27.6** (**30.69**) | **0.0180** | 0.08 |
| | IL-10 | 22.08 (18.51) | 17.89 (11.4) | 0.10 | 0.25 |
| | **IL-12p70** | **5.02** (**3.23**) | **4.17 (2.5)** | **0.0137** | 0.08 |
| | IL-13 | 10.04 (13.43) | 6.64 (11.56) | 0.08 | 0.25 |
| Monocyte activation markers (ng/mL) | CD14 | 2170.58 (872.73) | 2096.79 (884.5) | 0.83 | 0.83 |
| | CD163 | 626.82 (318.06) | 674.68 (519.74) | 0.14 | 0.25 |
| Neuroinflammatory markers (ng/mL) | NGAL | 159.46 (168.04) | 156.39 (119.54) | 0.63 | 0.70 |
| | MMP-9 | 801.98 (600.32) | 775.28 (647.17) | 0.83 | 0.83 |
| | YKL-40 | 27.87 (20.15) | 29.58 (30.97) | 0.10 | 0.17 |

Group data is presented as raw median (IQR). Statistical tests were conducted on log-scaled data. Analyses were corrected for multiple comparisons using the Benjamini-Hochberg procedure.

#### 1.4. Child serum markers at age 3 years

| Marker type | Marker | CHU (n=88) | CHEU (n=53) | P value Crude | Corrected |
|---|---|---|---|---|---|
| Pro-inflammatory cytokines (pg/mL) | **GM-CSF** | **66.75** (**95.11**) | **48.74** (**70.62**) | **0.0211** | 0.19 |
| | IFNγ | 18.94 (13.55) | 20.8 (16.48) | 0.24 | 0.50 |
| | IL-1β | 2.78 (1.99) | 2.81 (1.67) | 0.94 | 0.94 |
| | IL-2 | 2.62 (1.46) | 2.62 (2.1) | 0.56 | 0.81 |
| | IL-5 | 3.14 (3.47) | 2.79 (2.42) | 0.06 | 0.37 |
| | IL-6 | 4.9 (11.33) | 4.14 (6.84) | 0.84 | 0.94 |
| | IL-7 | 11.67 (5.94) | 12.7 (7.2) | 0.25 | 0.50 |
| | IL-8 | 18.65 (39.82) | 18.05 (23.7) | 0.0 | 0.95 |
| | TNFα | 12.76 (6.08) | 12.28 (7.62) | 0.66 | 0.91 |
| Anti-inflammatory cytokines (pg/mL) | IL-4 | 49.32 (106.84) | 39.95 (60.22) | 0.58 | 0.81 |
| | IL-10 | 10.49 (7.69) | 9.38 (5.22) | 0.11 | 0.40 |
| | IL-12p70 | 3.58 (2.31) | 4.12 (2.88) | 0.14 | 0.43 |
| | IL-13 | 11.6 (24.9) | 11.72 (19.1) | 0.78 | 0.93 |
| Monocyte activation markers (ng/mL) | **CD14** | **2255.06** (**990.69**) | **2538.94** (**922.53**) | **0.0135** | 0.24 |
| | CD163 | 661.71 (330.35) | 589.32 (334.17) | 0.13 | 0.43 |
| Neuroinflammatory markers (ng/mL) | NGAL | 115.62 (65.2) | 107.16 (58.34) | 0.21 | 0.53 |
| | MMP-9 | 901.36 (864.1) | 910.74 (1573.06) | 0.33 | 0.67 |
| | YKL-40 | 27.13 (20.31) | 26.33 (24.76) | 0.95 | 0.95 |

Group data is presented as raw median (IQR). Statistical tests were conducted on log-scaled data. Analyses were corrected for multiple comparisons using the Benjamini-Hochberg procedure.

#### **1.5. Child serum markers** at age 5 years

| **Marker type** | **Marker** | **CHU** (n=101) | **CHEU** (n=60) | **P value** Crude | Corrected |
|---|---|---|---|---|---|
| Pro-inflammatory cytokines (pg/mL) | GM-CSF | 76.78 (84.96) | 53.37 (66.61) | 0.07 | 0.27 |
| | IFNγ | 16.3 (9.7) | 17.8 (8.26) | 0.44 | 0.58 |
| | IL-1β | 2.77 (1.69) | 2.4 (1.56) | 0.43 | 0.61 |
| | IL-2 | 2.42 (1.83) | 2.12 (1.74) | 0.20 | 0.50 |
| | **IL-5** | **3.22** (**2.73**) | **2.3** (**2.21**) | **0.0102** | 0.09 |
| | IL-6 | 3.15 (7.48) | 2.08 (4.27) | 0.13 | 0.48 |
| | IL-7 | 10.28 (7.5) | 11.99 (8.4) | 0.65 | 0.73 |
| | IL-8 | 12.7 (32.83) | 18.15 (42.38) | 0.25 | 0.50 |
| | TNFα | 9.9 (5.34) | 9.51 (8.65) | 0.76 | 0.76 |
| Anti-inflammatory cytokines (pg/mL) | IL-4 | 36.18 (109.09) | 29.01 (41.65) | 0.28 | 0.58 |
| | **IL-10** | **9.13** (**7.00**) | **6.36** (**5.35**) | **0.0024** | **0.0439** |
| | IL-12p70 | 3.44 (2.32) | 3.22 (1.99) | 0.44 | 0.61 |
| | IL-13 | 7.88 (16.93) | 6.54 (14.78) | 0.65 | 0.73 |
| Monocyte activation markers (ng/mL) | CD14 | 2049.88 (1084.87) | 2295.11 (1009.77) | 0.13 | 0.40 |
| | CD163 | 523.98 (316.92) | 495.54 (235.2) | 0.70 | 0.74 |
| Neuroinflammatory markers (ng/mL) | NGAL | 105.76 (57.97) | 112.56 (59.23) | 0.35 | 0.61 |
| | **MMP-9** | **861.9** (**803.59**) | **1021.69** (**1264.14**) | **0.0471** | 0.27 |
| | YKL-40 | 19.92 (17.27) | 22.68 (18.24) | 0.49 | 0.62 |

Group data is presented as raw median (IQR). Statistical tests were conducted on log-scaled data. Analyses were corrected for multiple comparisons using the Benjamini-Hochberg procedure.

### Supplementary Table 2. Linear mixed-effects models

This table presents results from linear mixed-effects model examining child serum marker trajectories from birth to 5 years in children who are HIV-exposed uninfected (**CHEU**) and their HIV-unexposed (**CHU**) peers. Beta coefficients (**β**), standard errors (**SE**), t-values, degrees of freedom (**DF**), p-values, and Benjamini–Hochberg adjusted p-values (**BH p-values**) are reported for each orthogonal polynomial contrast (**OPC**): baseline (6 weeks), linear, quadratic, and cubic trajectories. **N obs** indicates the number of observations, **N sub** the number of subjects, and **ICC** the intraclass correlation coefficient from the model.

CHU are the reference group; estimates in CHU sections represent serum marker trends over time in these children. CHEU rows present the difference in serum marker trajectories relative to CHU at each corresponding OPC. A significant β at baseline for CHEU ($p<0.05$) indicates significantly different serum marker levels at birth compared to CHU. Significant interaction terms indicate that the longitudinal trajectory of a certain marker over time is different between groups; differences can be linear, quadratic, or cubic based on trajectory shapes.

| Marker type | Marker | Group | OPC | β | SE | T-value | DF | P-value | BH p-value | N obs | N sub | ICC |
|---|---|---|---|---|---|---|---|---|---|---|---|---|
| Pro-inflammatory | GM-CSF | CHU | Baseline | 4.009 | 0.057 | 70.59 | 206.6945 | **<0.001** | **0.003** | 623 | 215 | 0.295 |
| | | | Linear | 1.022 | 0.076 | 13.39 | 473.2402 | **<0.001** | **0.003** | | | |
| | | | Quadratic | -1.105 | 0.076 | -14.49 | 459.1556 | **<0.001** | **0.003** | | | |
| | | | Cubic | 0.684 | 0.077 | 8.92 | 457.3599 | **<0.001** | **0.003** | | | |
| | | CHEU | Baseline | -0.259 | 0.093 | -2.80 | 215.4157 | **0.006** | **0.015** | | | |
| | | | Linear | 0.015 | 0.127 | 0.12 | 485.4109 | 0.906 | 0.932 | | | |
| | | | Quadratic | 0.022 | 0.127 | 0.17 | 473.4586 | 0.865 | 0.909 | | | |
| | | | Cubic | 0.103 | 0.127 | 0.81 | 460.8547 | 0.417 | 0.561 | | | |
| | IFNγ | CHU | Baseline | 2.415 | 0.041 | 58.58 | 177.6853 | **<0.001** | **0.003** | 624 | 215 | 0.113 |
| | | | Linear | 0.966 | 0.070 | 13.84 | 476.9073 | **<0.001** | **0.003** | | | |
| | | | Quadratic | -0.467 | 0.070 | -6.66 | 460.8962 | **<0.001** | **0.003** | | | |
| | | | Cubic | -0.276 | 0.071 | -3.91 | 457.8532 | **<0.001** | **0.003** | | | |
| | | CHEU | Baseline | -0.035 | 0.068 | -0.51 | 188.2939 | 0.610 | 0.739 | | | |
| | | | Linear | 0.227 | 0.116 | 1.96 | 490.0138 | 0.050 | 0.101 | | | |
| | | | Quadratic | -0.005 | 0.116 | -0.04 | 477.1394 | 0.966 | 0.973 | | | |
| | | | Cubic | -0.124 | 0.117 | -1.06 | 462.0826 | 0.288 | 0.419 | | | |
| | IL-1β | CHU | Baseline | 0.660 | 0.044 | 14.92 | 189.1803 | **<0.001** | **0.003** | 623 | 215 | 0.212 |
| | | | Linear | 0.742 | 0.066 | 11.23 | 469.7764 | **<0.001** | **0.003** | | | |
| | | | Quadratic | -0.288 | 0.066 | -4.36 | 455.1048 | **<0.001** | **0.003** | | | |
| | | | Cubic | -0.060 | 0.067 | -0.90 | 452.1019 | 0.368 | 0.500 | | | |
| | | CHEU | Baseline | -0.156 | 0.072 | -2.16 | 198.4475 | **0.032** | 0.070 | | | |
| | | | Linear | 0.268 | 0.110 | 2.45 | 482.8661 | **0.015** | **0.035** | | | |
| | | | Quadratic | -0.043 | 0.110 | -0.39 | 470.2656 | 0.697 | 0.823 | | | |
| | | | Cubic | -0.100 | 0.110 | -0.91 | 455.9042 | 0.363 | 0.500 | | | |

| Marker type | Marker | Group | OPC | β | SE | T-value | DF | P-value | BH p-value | N obs | N sub | ICC |
|---|---|---|---|---|---|---|---|---|---|---|---|---|
| | IL-2 | CHU | Baseline | 0.651 | 0.041 | 15.72 | 186.8684 | **<0.001** | **0.003** | 624 | 215 | 0.128 |
| | | | Linear | 0.530 | 0.069 | 7.72 | 482.0314 | **<0.001** | **0.003** | | | |
| | | | Quadratic | -0.479 | 0.069 | -6.94 | 466.7276 | **<0.001** | **0.003** | | | |
| | | | Cubic | 0.207 | 0.070 | 2.97 | 463.8681 | **0.003** | **0.008** | | | |
| | | CHEU | Baseline | -0.100 | 0.068 | -1.47 | 197.5510 | 0.144 | 0.244 | | | |
| | | | Linear | 0.033 | 0.114 | 0.29 | 494.8908 | 0.773 | 0.870 | | | |
| | | | Quadratic | -0.027 | 0.114 | -0.24 | 482.4373 | 0.813 | 0.887 | | | |
| | | | Cubic | -0.216 | 0.115 | -1.88 | 467.9056 | 0.061 | 0.119 | | | |
| | IL-5 | CHU | Baseline | 0.958 | 0.050 | 19.00 | 201.3707 | **<0.001** | **0.003** | 624 | 215 | 0.249 |
| | | | Linear | 0.600 | 0.072 | 8.36 | 474.6514 | **<0.001** | **0.003** | | | |
| | | | Quadratic | -0.589 | 0.072 | -8.19 | 461.6827 | **<0.001** | **0.003** | | | |
| | | | Cubic | 0.180 | 0.072 | 2.48 | 459.6243 | **0.013** | **0.031** | | | |
| | | CHEU | Baseline | -0.114 | 0.083 | -1.38 | 210.8371 | 0.169 | 0.283 | | | |
| | | | Linear | -0.439 | 0.119 | -3.68 | 488.1433 | **<0.001** | **0.003** | | | |
| | | | Quadratic | 0.156 | 0.120 | 1.30 | 476.4356 | 0.193 | 0.312 | | | |
| | | | Cubic | -0.088 | 0.120 | -0.74 | 463.1634 | 0.461 | 0.603 | | | |
| | IL-6 | CHU | Baseline | 1.038 | 0.080 | 12.96 | 197.8931 | **<0.001** | **0.003** | 624 | 215 | 0.202 |
| | | | Linear | 0.520 | 0.121 | 4.30 | 478.9070 | **<0.001** | **0.003** | | | |
| | | | Quadratic | -0.591 | 0.121 | -4.88 | 465.0803 | **<0.001** | **0.003** | | | |
| | | | Cubic | -0.063 | 0.122 | -0.52 | 462.7333 | 0.605 | 0.739 | | | |
| | | CHEU | Baseline | -0.206 | 0.131 | -1.57 | 207.8671 | 0.118 | 0.207 | | | |
| | | | Linear | -0.197 | 0.201 | -0.98 | 492.1695 | 0.326 | 0.465 | | | |
| | | | Quadratic | -0.241 | 0.201 | -1.20 | 480.1668 | 0.232 | 0.355 | | | |
| | | | Cubic | -0.147 | 0.202 | -0.73 | 466.4200 | 0.466 | 0.604 | | | |
| | IL-7 | CHU | Baseline | 2.137 | 0.034 | 62.35 | 183.4732 | **<0.001** | **0.003** | 624 | 215 | 0.116 |
| | | | Linear | 0.326 | 0.058 | 5.65 | 481.4469 | **<0.001** | **0.003** | | | |
| | | | Quadratic | -0.403 | 0.058 | -6.94 | 465.8221 | **<0.001** | **0.003** | | | |
| | | | Cubic | 0.021 | 0.058 | 0.36 | 462.8580 | 0.721 | 0.842 | | | |
| | | CHEU | Baseline | -0.005 | 0.056 | -0.08 | 194.2376 | 0.935 | 0.955 | | | |
| | | | Linear | 0.018 | 0.096 | 0.19 | 494.2747 | 0.849 | 0.899 | | | |
| | | | Quadratic | 0.043 | 0.096 | 0.44 | 481.7045 | 0.657 | 0.782 | | | |
| | | | Cubic | -0.149 | 0.097 | -1.54 | 466.9879 | 0.125 | 0.214 | | | |
| | IL-8 | CHU | Baseline | 2.589 | 0.074 | 35.17 | 201.9011 | **<0.001** | **0.003** | 624 | 215 | 0.244 |
| | | | Linear | 0.556 | 0.105 | 5.28 | 475.7419 | **<0.001** | **0.003** | | | |
| | | | Quadratic | -0.227 | 0.105 | -2.16 | 462.7308 | **0.032** | 0.070 | | | |
| | | | Cubic | -0.141 | 0.106 | -1.33 | 460.6519 | 0.184 | 0.301 | | | |
| | | CHEU | Baseline | 0.026 | 0.120 | 0.22 | 211.4342 | 0.830 | 0.892 | | | |
| | | | Linear | 0.188 | 0.175 | 1.07 | 489.1732 | 0.284 | 0.417 | | | |
| | | | Quadratic | 0.216 | 0.175 | 1.23 | 477.4727 | 0.220 | 0.344 | | | |
| | | | Cubic | -0.062 | 0.176 | -0.35 | 464.1936 | 0.725 | 0.842 | | | |
| | TNFα | CHU | Baseline | 2.572 | 0.036 | 71.87 | 169.3344 | **<0.001** | **0.003** | 624 | 215 | 0.105 |
| | | | Linear | -0.382 | 0.061 | -6.24 | 470.9187 | **<0.001** | **0.003** | | | |
| | | | Quadratic | 0.135 | 0.062 | 2.19 | 454.2980 | **0.029** | 0.065 | | | |
| | | | Cubic | -0.097 | 0.062 | -1.57 | 451.1126 | 0.117 | 0.207 | | | |

| Marker type | Marker | Group | OPC | β | SE | T-value | DF | P-value | BH p-value | N obs | N sub | ICC |
|---|---|---|---|---|---|---|---|---|---|---|---|---|
| | | CHEU | Baseline | -0.009 | 0.059 | -0.15 | 179.7699 | 0.882 | 0.920 | | | |
| | | | Linear | 0.191 | 0.101 | 1.88 | 484.3450 | 0.061 | 0.119 | | | |
| | | | Quadratic | 0.035 | 0.102 | 0.34 | 471.0527 | 0.731 | 0.842 | | | |
| | | | Cubic | -0.063 | 0.103 | -0.62 | 455.5061 | 0.539 | 0.675 | | | |
| Anti-inflammatory | IL-4 | CHU | Baseline | 3.425 | 0.077 | 44.72 | 198.8740 | **<0.001** | **0.003** | 624 | 215 | 0.245 |
| | | | Linear | 0.780 | 0.109 | 7.12 | 473.1659 | **<0.001** | **0.003** | | | |
| | | | Quadratic | -0.623 | 0.110 | -5.68 | 460.0141 | **<0.001** | **0.003** | | | |
| | | | Cubic | -0.078 | 0.110 | -0.71 | 457.9148 | 0.479 | 0.610 | | | |
| | | CHEU | Baseline | -0.212 | 0.125 | -1.69 | 208.3310 | 0.092 | 0.170 | | | |
| | | | Linear | 0.024 | 0.182 | 0.13 | 486.7626 | 0.893 | 0.925 | | | |
| | | | Quadratic | -0.044 | 0.182 | -0.24 | 474.9205 | 0.810 | 0.887 | | | |
| | | | Cubic | -0.107 | 0.183 | -0.58 | 461.4938 | 0.559 | 0.694 | | | |
| | IL-10 | CHU | Baseline | 2.484 | 0.042 | 59.68 | 187.1844 | **<0.001** | **0.003** | 624 | 215 | 0.193 |
| | | | Linear | -0.418 | 0.063 | -6.59 | 471.1790 | **<0.001** | **0.003** | | | |
| | | | Quadratic | -0.390 | 0.064 | -6.13 | 456.6628 | **<0.001** | **0.003** | | | |
| | | | Cubic | 0.259 | 0.064 | 4.04 | 454.1742 | **<0.001** | **0.003** | | | |
| | | CHEU | Baseline | -0.184 | 0.068 | -2.70 | 196.9789 | **0.008** | **0.020** | | | |
| | | | Linear | -0.187 | 0.105 | -1.77 | 484.9374 | 0.077 | 0.144 | | | |
| | | | Quadratic | -0.032 | 0.106 | -0.30 | 472.3858 | 0.763 | 0.865 | | | |
| | | | Cubic | -0.080 | 0.106 | -0.75 | 458.0261 | 0.451 | 0.596 | | | |
| | IL-12p70 | CHU | Baseline | 1.152 | 0.040 | 28.88 | 184.2989 | **<0.001** | **0.003** | 624 | 215 | 0.180 |
| | | | Linear | 0.262 | 0.062 | 4.24 | 470.8358 | **<0.001** | **0.003** | | | |
| | | | Quadratic | -0.421 | 0.062 | -6.79 | 455.9783 | **<0.001** | **0.003** | | | |
| | | | Cubic | 0.210 | 0.062 | 3.35 | 453.3863 | **<0.001** | **0.003** | | | |
| | | CHEU | Baseline | -0.072 | 0.065 | -1.10 | 194.1808 | 0.272 | 0.408 | | | |
| | | | Linear | 0.203 | 0.103 | 1.97 | 484.5984 | **0.049** | 0.101 | | | |
| | | | Quadratic | -0.066 | 0.103 | -0.64 | 471.8905 | 0.525 | 0.663 | | | |
| | | | Cubic | -0.164 | 0.103 | -1.59 | 457.3127 | 0.113 | 0.206 | | | |
| | IL-13 | CHU | Baseline | 1.919 | 0.089 | 21.55 | 204.4346 | **<0.001** | **0.003** | 624 | 215 | 0.319 |
| | | | Linear | 0.742 | 0.116 | 6.40 | 467.3485 | **<0.001** | **0.003** | | | |
| | | | Quadratic | -0.920 | 0.116 | -7.93 | 455.6742 | **<0.001** | **0.003** | | | |
| | | | Cubic | 0.235 | 0.117 | 2.01 | 454.0130 | **0.045** | 0.095 | | | |
| | | CHEU | Baseline | -0.132 | 0.145 | -0.91 | 213.1386 | 0.366 | 0.500 | | | |
| | | | Linear | -0.059 | 0.193 | -0.31 | 480.9932 | 0.758 | 0.865 | | | |
| | | | Quadratic | 0.140 | 0.193 | 0.72 | 469.8473 | 0.470 | 0.604 | | | |
| | | | Cubic | -0.247 | 0.193 | -1.28 | 457.3701 | 0.202 | 0.323 | | | |
| Monocyte activation | CD14 | CHU | Baseline | 7.584 | 0.020 | 370.91 | 191.0327 | **<0.001** | **0.003** | 625 | 215 | 0.237 |
| | | | Linear | 0.219 | 0.030 | 7.43 | 468.0744 | **<0.001** | **0.003** | | | |
| | | | Quadratic | -0.197 | 0.030 | -6.66 | 454.3850 | **<0.001** | **0.003** | | | |
| | | | Cubic | 0.047 | 0.030 | 1.57 | 452.1787 | 0.117 | 0.207 | | | |
| | | CHEU | Baseline | 0.086 | 0.033 | 2.58 | 200.0964 | **0.011** | **0.026** | | | |
| | | | Linear | -0.012 | 0.049 | -0.24 | 480.8668 | 0.810 | 0.887 | | | |
| | | | Quadratic | 0.045 | 0.049 | 0.92 | 468.6972 | 0.361 | 0.500 | | | |
| | | | Cubic | -0.125 | 0.049 | -2.54 | 457.2720 | **0.011** | **0.026** | | | |

| **Marker type** | **Marker** | **Group** | **OPC** | **β** | **SE** | **T-value** | **DF** | **P-value** | **BH p-value** | **N obs** | **N sub** | **ICC** |
|---|---|---|---|---|---|---|---|---|---|---|---|---|
| | CD163 | CHU | Baseline | 6.403 | 0.031 | 203.42 | 208.1133 | **<0.001** | **0.003** | 625 | 215 | 0.402 |
| | | | Linear | -0.068 | 0.037 | -1.84 | 460.4099 | 0.067 | 0.129 | | | |
| | | | Quadratic | -0.162 | 0.037 | -4.40 | 450.3383 | **<0.001** | **0.003** | | | |
| | | | Cubic | -0.029 | 0.037 | -0.79 | 449.1004 | 0.430 | 0.573 | | | |
| | | CHEU | Baseline | -0.013 | 0.051 | -0.26 | 215.5244 | 0.796 | 0.887 | | | |
| | | | Linear | -0.095 | 0.061 | -1.55 | 472.5353 | 0.121 | 0.210 | | | |
| | | | Quadratic | -0.004 | 0.061 | -0.06 | 462.5814 | 0.949 | 0.962 | | | |
| | | | Cubic | 0.120 | 0.061 | 1.96 | 453.2366 | 0.050 | 0.101 | | | |
| Neuroinflammatory | NGAL | CHU | Baseline | 4.787 | 0.029 | 166.89 | 188.4458 | **<0.001** | **0.003** | 624 | 215 | 0.156 |
| | | | Linear | 0.020 | 0.046 | 0.45 | 478.2307 | 0.655 | 0.782 | | | |
| | | | Quadratic | -0.317 | 0.046 | -6.88 | 463.3064 | **<0.001** | **0.003** | | | |
| | | | Cubic | 0.262 | 0.046 | 5.65 | 460.6166 | **<0.001** | **0.003** | | | |
| | | CHEU | Baseline | -0.064 | 0.047 | -1.36 | 199.4345 | 0.176 | 0.291 | | | |
| | | | Linear | 0.146 | 0.076 | 1.91 | 492.0003 | 0.056 | 0.112 | | | |
| | | | Quadratic | 0.018 | 0.076 | 0.23 | 479.0493 | 0.819 | 0.887 | | | |
| | | | Cubic | 0.080 | 0.077 | 1.04 | 466.0983 | 0.300 | 0.432 | | | |
| | MMP-9 | CHU | Baseline | 6.529 | 0.042 | 154.93 | 200.0863 | **<0.001** | **0.003** | 624 | 215 | 0.209 |
| | | | Linear | 0.502 | 0.063 | 7.97 | 479.3032 | **<0.001** | **0.003** | | | |
| | | | Quadratic | -0.346 | 0.063 | -5.48 | 465.6838 | **<0.001** | **0.003** | | | |
| | | | Cubic | 0.032 | 0.064 | 0.51 | 463.3946 | 0.611 | 0.739 | | | |
| | | CHEU | Baseline | 0.083 | 0.069 | 1.21 | 210.4898 | 0.228 | 0.353 | | | |
| | | | Linear | 0.186 | 0.105 | 1.77 | 492.8931 | 0.077 | 0.144 | | | |
| | | | Quadratic | 0.021 | 0.105 | 0.20 | 480.6255 | 0.843 | 0.899 | | | |
| | | | Cubic | -0.003 | 0.105 | -0.03 | 468.4308 | 0.977 | 0.977 | | | |
| | YKL-40 | CHU | Baseline | 3.297 | 0.039 | 83.48 | 190.6215 | **<0.001** | **0.003** | 625 | 215 | 0.396 |
| | | | Linear | -0.233 | 0.047 | -5.01 | 444.9912 | **<0.001** | **0.003** | | | |
| | | | Quadratic | -0.057 | 0.047 | -1.23 | 434.2399 | 0.218 | 0.344 | | | |
| | | | Cubic | -0.094 | 0.047 | -2.01 | 432.9085 | **0.045** | 0.095 | | | |
| | | CHEU | Baseline | 0.073 | 0.064 | 1.13 | 197.7634 | 0.258 | 0.391 | | | |
| | | | Linear | 0.071 | 0.078 | 0.92 | 457.8959 | 0.358 | 0.500 | | | |
| | | | Quadratic | -0.085 | 0.078 | -1.09 | 447.2502 | 0.275 | 0.408 | | | |
| | | | Cubic | 0.183 | 0.077 | 2.37 | 437.2982 | **0.018** | **0.041** | | | |

**Supplementary Table 3. Group differences in parietal brain metabolite levels at 6–7 years**

| Brain region | Neurometabolite | MRS measurement | CHU | CHEU | β | 95% CI | P-value |
|---|---|---|---|---|---|---|---|
| **Midline parietal grey matter** | **Glutamate** | **Ratios to creatine** | **1.68** (**0.23**) | **1.60** (**0.27**) | **0.10** | (**0.03 to 0.63**) | **0.0308** |
| | | Absolute concentrations | 11.98 (2.52) | 11.69 (2.55) | 0.07 | (-0.08 to 0.55) | 0.14 |
| | Myo-inositol | Ratios to creatine | 0.85 (0.07) | 0.84 (0.06) | 0.06 | (-0.08 to 0.50) | 0.16 |
| | | Absolute concentrations | 6.15 (1.22) | 6.02 (0.71) | 0.04 | (-0.15 to 0.43) | 0.35 |
| | N-acetyl-aspartate | Ratios to creatine | 1.38 (0.12) | 1.36 (0.1) | 0.02 | (-0.24 to 0.37) | 0.68 |
| | | Absolute concentrations | 10.15 (2.1) | 9.69 (1.31) | 0.02 | (-0.23 to 0.38) | 0.64 |
| | Total choline | Ratios to creatine | 0.17 (0.02) | 0.17 (0.02) | -0.01 | (-0.37 to 0.32) | 0.90 |
| | | Absolute concentrations | 1.24 (0.24) | 1.25 (0.25) | -0.02 | (-0.42 to 0.27) | 0.68 |
| | Total creatine | Absolute concentrations | 7.34 (1.15) | 7.06 (1.00) | 0.00 | (-0.33 to 0.33) | 0.99 |
| **Left parietal white matter** | Glutamate | Ratios to creatine | 1.41 (0.28) | 1.42 (0.3) | 0.03 | (-0.28 to 0.49) | 0.58 |
| | | Absolute concentrations | 10.03 (1.72) | 10.62 (3.15) | -0.06 | (-1.07 to 0.39) | 0.34 |
| | Myo-inositol | Ratios to creatine | 0.86 (0.11) | 0.87 (0.08) | -0.04 | (-0.45 to 0.17) | 0.39 |
| | | Absolute concentrations | 6.12 (1.07) | 6.4 (1.04) | -0.10 | (-1.11 to 0.04) | 0.0646 |
| | N-acetyl-aspartate | Ratios to creatine | 1.55 (0.15) | 1.52 (0.19) | 0.06 | (-0.16 to 0.61) | 0.24 |
| | | Absolute concentrations | 11.1 (1.84) | 11.31 (2.78) | 0.01 | (-0.66 to 0.75) | 0.90 |
| | **Total choline** | **Ratios to creatine** | **0.25** (**0.03**) | **0.23** (**0.04**) | **0.13** | (**0.04 to 0.82**) | **0.0295** |
| | | Absolute concentrations | 1.72 (0.28) | 1.66 (0.34) | 0.00 | (-0.72 to 0.67) | 0.94 |
| | Total creatine | Absolute concentrations | 7.01 (1.06) | 7.18 (0.81) | -0.03 | (-0.80 to 0.42) | 0.53 |

Group data is presented as raw median (IQR); statistical tests were conducted on scaled data. High-quality MRS data available in the midline parietal grey matter: n=131 CHU and n=57 CHEU (ratios to creatine); n=127 CHU and n=57 CHEU (absolute concentrations, in millimolar). In the left parietal white matter: n=116 CHU and n=47 CHEU (ratios to creatine); n=74 CHU and n=16 CHEU (absolute concentrations, in millimolar). **CHEU**: Children who are HIV-Exposed and Uninfected; **CHU**: Children who are HIV-Unexposed; **MRS**: Magnetic Resonance Spectroscopy; **β**: Effect size.

### Supplementary Table 4. Associations between maternal/child serum marker levels at different timepoints and child neurometabolite ratios to creatine at 6–7 years

A full Statistical Analysis Report, including all unadjusted and adjusted results, has been deposited on OSF and is publicly available under DOI: 10.17605/OSF.IO/7N94C.

***Maternal serum marker levels during pregnancy***

Glutamate: Maternal **IL-8** levels during pregnancy were negatively associated with glutamate ratios in the left parietal white matter of children who are HEU (β=−0.73, 95% CI −1.13 to −0.33, p=0.0004). No significant associations were observed in HU peers.

Total choline: In the midline parietal grey matter, maternal **IL-6** and **IL-4** were positively associated with total choline ratios in HU children (IL-6: β=0.25, 95% CI 0.10 to 0.40, p=0.0017; IL-4: β=0.21, 95% CI 0.06 to 0.37, p=0.0077). Maternal HIV significantly altered these relationships, and no significant associations were detected in children who are HEU.

N-acetyl-aspartate: Maternal **IL-8** was negatively associated with lower N-acetyl-aspartate ratios in the left parietal white matter of HEU children only (β=−0.69, 95% CI −1.22 to −0.17, p=0.0101).

***Child serum marker trajectories from 6 weeks to 5 years***

Myo-inositol: Slopes of **IL-8**, **TNFα**, and **IL-4** from infancy to 5 years were negatively associated with myo-inositol ratios in the midline parietal grey matter of HU children (IL-8: β=−0.28, 95% CI −0.49 to −0.06, p=0.0124; TNFα: β=−0.44, 95% CI −0.72 to −0.16, p=0.0023; IL-4: β=−0.29, 95% CI −0.49 to −0.09, p=0.0048). No significant associations were observed in HEU peers.

Total choline: In children who are HEU, **IL-8** trajectories were negatively associated with total choline ratios in the midline parietal grey matter (β=−0.34, 95% CI −0.68 to −0.01, p=0.0461). In HU children, TNFα trajectories were negatively associated with total choline ratios in the left parietal white matter (β=−0.32, 95% CI −0.54 to −0.09, p=0.0060).

***Week 6 serum marker levels***

Myo-inositol: **sCD14** levels during infancy were negatively associated with lower myo-inositol ratios in the midline parietal grey matter of HU children (β=−1.43, 95% CI −2.53 to −0.34, p=0.0107). No significant associations were observed in HEU peers.

***Year 2 serum marker levels***

Glutamate: In HU children, **IL-13** was positively associated with left parietal white matter glutamate ratios (β=0.28, 95% CI 0.08 to 0.47, p=0.0058). No significant associations were observed in HEU peers.

***Year 3 serum marker levels***

Glutamate: At age 3 years, **IL-4** and **IL-13** were negatively associated with glutamate ratios in the left parietal white matter of children who are HEU (IL-4: β=−0.87, 95% CI −1.36 to −0.37, p=0.0007; IL-13: β=−0.67, 95% CI −1.06 to −0.28, p=0.0009). No significant associations were observed in HU peers.

***Year 5 serum marker levels***

Glutamate: At age 5 years, **IL-8** was negatively associated with glutamate ratios in the midline parietal grey matter of HU children (β=−0.21, 95% CI −0.36 to −0.05, p=0.0100). No significant associations were observed in HEU peers.

Myo-inositol: Several markers measured at age 5 years were negatively associated with myo-inositol ratios in HU children: **IL-8** (β=−0.19, 95% CI −0.35 to −0.04, p=0.0143), **TNFα** (β=−0.44, 95% CI −0.71 to −0.16, p=0.0022), **IL-4** (β=−0.22, 95% CI −0.40 to −0.04, p=0.0192), **IL-13** (β=−0.20, 95% CI −0.35 to −0.04, p=0.0164), and **NGAL** (β=−0.66, 95% CI −1.25 to −0.07, p=0.0298). Only **IL-4** showed a significant association in children who are HEU (β=0.22, 95% CI 0.01 to 0.44, p=0.0399).

Total choline: In HEU children, **IL-1β** was negatively associated with total choline ratios in the midline parietal grey matter (β=−0.63, 95% CI −1.16 to −0.09, p=0.0217). No significant associations were observed in HU peers.

**Supplementary Table 5. Adjusted linear models exploring associations between maternal/child serum marker levels at different timepoints and midline parietal grey matter neurometabolite absolute concentrations at 6–7 years**

| | | | | | | | CHU | | CHEU | |
|---|---|---|---|---|---|---|---|---|---|---|
| **Serum** | **Timepoint** | **n CHU** | **n CHEU** | **Neurometabolite** | **Marker type** | **Marker** | **β** (95% CI) | **P-value** | **β** (95% CI) | **P-value** |
| Child | 6 weeks | 70 | 40 | Choline | Anti-inflammatory | IL-4 | – | – | –0.27 (–0.51 to –0.03) | **0.0310** |
| | 2 years | 76 | 39 | Glutamate | Monocyte activation | CD14 | –0.86 (–1.54 to –0.18) | **0.0135** | – | – |
| | | | | Myo-inositol | Pro-inflammatory | TNFα | – | – | –0.47 (–0.92 to –0.02) | **0.0422** |
| | | | | | Monocyte activation | CD14 | –0.74 (–1.43 to –0.06) | **0.0333** | – | – |
| | 3 years | 69 | 38 | Glutamate | Pro-inflammatory | IFNγ | –0.38 (–0.71 to –0.04) | **0.0288** | – | – |
| | | | | | Monocyte activation | CD14 | –0.71 (–1.41 to –0.01) | **0.0495** | – | – |
| | | | | N-acetyl-aspartate | Pro-inflammatory | IL-5 | 0.28 (0.03 to 0.54) | **0.0321** | – | – |

Linear regression models with robust standard errors. Models were adjusted for child age, child sex, and voxel tissue composition. **CHEU**: Children who are HIV-Exposed and Uninfected; **CHU**: Children who are HIV-Unexposed; **β**: Effect size.

Please note: Association analyses using child absolute neurometabolite concentrations in the left parietal white matter are not presented because the small sample size in this region (CHEU n=16) precluded adequately adjusted models and yielded unstable unadjusted estimates. As prespecified, only adjusted, BH-corrected associations are reported.

### Supplementary Table 6. Sensitivity analyses

#### 6.1. Maternal age at delivery

| | | | | | | | | CHU | | CHEU | |
|---|---|---|---|---|---|---|---|---|---|---|---|
| **Serum** | **Timepoint** | **Brain region** | **n CHU** | **n CHEU** | **Neurometabolite** | **Marker type** | **Marker** | **β** (95% CI) | **P-value** | **β** (95% CI) | **P-value** |
| Maternal | Pregnancy | GM | 99 | 54 | Choline | Pro-inflammatory | IL-6 | 0.25 (0.10 to 0.41) | **0.0017** | – | – |
| | | | | | | Anti-inflammatory | IL-4 | 0.21 (0.06 to 0.37) | **0.0070** | – | – |
| | | WM | 88 | 44 | Glutamate | Pro-inflammatory | IL-8 | – | – | –0.75 (–1.13 to –0.36) | **0.0002** |
| | | | | | N-acetyl-aspartate | Pro-inflammatory | IL-8 | – | – | –0.69 (–1.22 to –0.15) | **0.0131** |
| Child | 6 weeks | GM | 75 | 40 | Myo-inositol | Monocyte activation | CD14 | –1.37 (–2.44 to –0.29) | **0.0135** | – | – |
| | 2 years | WM | 72 | 32 | Glutamate | Anti-inflammatory | IL-13 | 0.26 (0.06 to 0.46) | **0.0058** | – | – |
| | 3 years | WM | 62 | 31 | Glutamate | Anti-inflammatory | IL-4 | – | – | –0.87 (–1.36 to –0.37) | **0.0008** |
| | | | | | | | IL-13 | – | – | –0.67 (–1.07 to –0.28) | **0.0010** |
| | 5 years | GM | 81 | 45 | Glutamate | Pro-inflammatory | IL-8 | –0.21 (–0.37 to –0.05) | **0.0116** | – | – |
| | | | | | Myo-inositol | Pro-inflammatory | IL-8 | –0.20 (–0.35 to –0.05) | **0.0095** | – | – |
| | | | | | | | TNFα | –0.48 (–0.76 to –0.20) | **0.0008** | – | – |
| | | | | | | Anti-inflammatory | IL-4 | –0.24 (–0.42 to –0.06) | **0.0098** | 0.25 (0.03 to 0.46) | **0.0235** |
| | | | | | | | IL-13 | –0.22 (–0.38 to –0.06) | **0.0072** | – | – |
| | | | | | | Neuroinflammatory | NGAL | –0.69 (–1.28 to –0.10) | **0.0222** | – | – |
| | | | | | Choline | Pro-inflammatory | IL-1β | – | – | –0.68 (–1.21 to –0.14) | **0.0134** |

Linear regression models with robust standard errors. Adjusted for child age, child sex, voxel composition, and **maternal age at delivery**.

### 6.2. Maternal alcohol use during pregnancy

| | | | | | | | | CHU | | CHEU | |
|---|---|---|---|---|---|---|---|---|---|---|---|
| **Serum** | **Timepoint** | **Brain region** | **n CHU** | **n CHEU** | **Neurometabolite** | **Marker type** | **Marker** | **β** (95% CI) | **P-value** | **β** (95% CI) | **P-value** |
| Maternal | Pregnancy | GM | 99 | 54 | Choline | Pro-inflammatory | IL-6 | 0.25 (0.09 to 0.40) | **0.0020** | – | – |
| | | | | | | Anti-inflammatory | IL-4 | 0.21 (0.05 to 0.36) | **0.0092** | – | – |
| | | WM | 88 | 44 | Glutamate | Pro-inflammatory | IL-8 | – | – | –0.76 (–1.20 to –0.31) | **0.0010** |
| | | | | | N-acetyl-aspartate | Pro-inflammatory | IL-8 | – | – | –0.86 (–1.31 to –0.41) | **0.0002** |
| Child | 6 weeks | GM | 75 | 40 | Myo-inositol | Monocyte activation | CD14 | –1.44 (–2.54 to –0.34) | **0.0105** | – | – |
| | 2 years | WM | 72 | 32 | Glutamate | Anti-inflammatory | IL-13 | 0.29 (0.07 to 0.50) | **0.0096** | – | – |
| | 3 years | WM | 62 | 31 | Glutamate | Anti-inflammatory | IL-4 | – | – | –0.84 (–1.40 to –0.27) | **0.0043** |
| | | | | | | | IL-13 | – | – | –0.65 (–1.10 to –0.21) | **0.0048** |
| | 5 years | GM | 81 | 45 | Glutamate | Pro-inflammatory | IL-8 | –0.19 (–0.34 to –0.04) | **0.0139** | – | – |
| | | | | | Myo-inositol | Pro-inflammatory | IL-8 | –0.18 (–0.33 to –0.03) | **0.0211** | – | – |
| | | | | | | | TNFα | –0.41 (–0.68 to –0.14) | **0.0036** | – | – |
| | | | | | | Anti-inflammatory | IL-4 | –0.21 (–0.38 to –0.04) | **0.0178** | 0.18 (–0.03 to 0.40) | 0.09 |
| | | | | | | | IL-13 | –0.20 (–0.35 to –0.05) | **0.0115** | – | – |
| | | | | | | Neuroinflammatory | NGAL | –0.62 (–1.23 to –0.01) | **0.0449** | – | – |
| | | | | | Choline | Pro-inflammatory | IL-1β | – | – | –0.79 (–1.36 to –0.22) | **0.0067** |

Linear regression models with robust standard errors. Adjusted for child age, child sex, voxel composition, and **maternal alcohol use during pregnancy**.

### 6.3. Maternal depression during pregnancy

| | | | | | | | | CHU | | CHEU | |
|---|---|---|---|---|---|---|---|---|---|---|---|
| **Serum** | **Timepoint** | **Brain region** | **n CHU** | **n CHEU** | **Neurometabolite** | **Marker type** | **Marker** | **β** (95% CI) | **P-value** | **β** (95% CI) | **P-value** |
| Maternal | Pregnancy | GM | 99 | 54 | Choline | Pro-inflammatory | IL-6 | 0.21 (0.07 to 0.36) | **0.0052** | – | – |
| | | | | | | Anti-inflammatory | IL-4 | 0.18 (0.02 to 0.33) | **0.0284** | – | – |
| | | WM | 88 | 44 | Glutamate | Pro-inflammatory | IL-8 | – | – | –0.74 (–1.15 to –0.33) | **0.0005** |
| | | | | | N-acetyl-aspartate | Pro-inflammatory | IL-8 | – | – | –0.70 (–1.23 to –0.17) | **0.0100** |
| Child | 6 weeks | GM | 75 | 40 | Myo-inositol | Monocyte activation | CD14 | –1.41 (–2.50 to –0.33) | **0.0112** | – | – |
| | 2 years | WM | 72 | 32 | Glutamate | Anti-inflammatory | IL-13 | 0.28 (0.08 to 0.47) | **0.0064** | – | – |
| | 3 years | WM | 62 | 31 | Glutamate | Anti-inflammatory | IL-4 | – | – | –0.86 (–1.36 to –0.36) | **0.0010** |
| | | | | | | | IL-13 | – | – | –0.67 (–1.08 to –0.27) | **0.0013** |
| | 5 years | GM | 81 | 45 | Glutamate | Pro-inflammatory | IL-8 | –0.20 (–0.38 to –0.03) | **0.0228** | – | – |
| | | | | | Myo-inositol | Pro-inflammatory | IL-8 | –0.22 (–0.39 to –0.05) | **0.0121** | – | – |
| | | | | | | | TNFα | –0.45 (–0.73 to –0.16) | **0.0023** | – | – |
| | | | | | | Anti-inflammatory | IL-4 | –0.24 (–0.42 to –0.06) | **0.0114** | 0.23 (0.02 to 0.45) | **0.0349** |
| | | | | | | | IL-13 | –0.22 (–0.39 to –0.05) | **0.0113** | – | – |
| | | | | | | Neuroinflammatory | NGAL | –0.69 (–1.34 to –0.05) | **0.0362** | – | – |
| | | | | | Choline | Pro-inflammatory | IL-1β | – | – | –0.65 (–1.19 to –0.11) | **0.0184** |

Linear regression models with robust standard errors. Adjusted for child age, child sex, voxel composition, and **maternal depression during pregnancy**.

### 6.4. Maternal smoking during pregnancy

| | | | | | | | | CHU | | CHEU | |
|---|---|---|---|---|---|---|---|---|---|---|---|
| **Serum** | **Timepoint** | **Brain region** | **n CHU** | **n CHEU** | **Neurometabolite** | **Marker type** | **Marker** | **β** (95% CI) | **P-value** | **β** (95% CI) | **P-value** |
| Maternal | Pregnancy | GM | 99 | 54 | Choline | Pro-inflammatory | IL-6 | 0.23 (0.08 to 0.38) | **0.0030** | – | – |
| | | | | | | Anti-inflammatory | IL-4 | 0.21 (0.07 to 0.35) | **0.0041** | – | – |
| | | WM | 88 | 44 | Glutamate | Pro-inflammatory | IL-8 | – | – | –0.74 (–1.16 to –0.33) | **0.0005** |
| | | | | | N-acetyl-aspartate | Pro-inflammatory | IL-8 | – | – | –0.70 (–1.22 to –0.18) | **0.0092** |
| Child | 6 weeks | GM | 75 | 40 | Myo-inositol | Monocyte activation | CD14 | –1.52 (–2.59 to –0.45) | **0.0059** | – | – |
| | 2 years | WM | 72 | 32 | Glutamate | Anti-inflammatory | IL-13 | 0.27 (0.09 to 0.46) | **0.0045** | – | – |
| | 3 years | WM | 62 | 31 | Glutamate | Anti-inflammatory | IL-4 | – | – | –0.89 (–1.38 to –0.40) | **0.0006** |
| | | | | | | | IL-13 | – | – | –0.69 (–1.07 to –0.31) | **0.0006** |
| | 5 years | GM | 81 | 45 | Glutamate | Pro-inflammatory | IL-8 | –0.21 (–0.36 to –0.05) | **0.0117** | – | – |
| | | | | | Myo-inositol | Pro-inflammatory | IL-8 | –0.19 (–0.35 to –0.04) | **0.0161** | – | – |
| | | | | | | | TNFα | –0.44 (–0.72 to –0.16) | **0.0025** | – | – |
| | | | | | | Anti-inflammatory | IL-4 | –0.22 (–0.40 to –0.04) | **0.0161** | 0.23 (0.01 to 0.45) | **0.0398** |
| | | | | | | | IL-13 | –0.20 (–0.36 to –0.04) | **0.0146** | – | – |
| | | | | | | Neuroinflammatory | NGAL | –0.68 (–1.31 to –0.05) | **0.0351** | – | – |
| | | | | | Choline | Pro-inflammatory | IL-1β | – | – | –0.61 (–1.15 to –0.07) | **0.0268** |

Linear regression models with robust standard errors. Adjusted for child age, child sex, voxel composition, and **maternal smoking during pregnancy**.

### 6.5. Exclusive breastfeeding duration

| | | | | | | | | CHU | | CHEU | |
|---|---|---|---|---|---|---|---|---|---|---|---|
| **Serum** | **Timepoint** | **Brain region** | **n CHU** | **n CHEU** | **Neurometabolite** | **Marker type** | **Marker** | **β** (95% CI) | **P-value** | **β** (95% CI) | **P-value** |
| Maternal | Pregnancy | GM | 99 | 54 | Choline | Pro-inflammatory | IL-6 | 0.25 (0.10 to 0.41) | **0.0015** | – | – |
| | | | | | | Anti-inflammatory | IL-4 | 0.22 (0.06 to 0.38) | **0.0068** | – | – |
| | | WM | 88 | 44 | Glutamate | Pro-inflammatory | IL-8 | – | – | –0.74 (–1.15 to –0.32) | **0.0006** |
| | | | | | N-acetyl-aspartate | Pro-inflammatory | IL-8 | – | – | –0.69 (–1.21 to –0.17) | **0.0097** |
| Child | 6 weeks | GM | 75 | 40 | Myo-inositol | Monocyte activation | CD14 | –1.46 (–2.61 to –0.31) | **0.0137** | – | – |
| | 2 years | WM | 72 | 32 | Glutamate | Anti-inflammatory | IL-13 | 0.28 (0.08 to 0.47) | **0.0061** | – | – |
| | 3 years | WM | 62 | 31 | Glutamate | Anti-inflammatory | IL-4 | – | – | –0.88 (–1.42 to –0.34) | **0.0016** |
| | | | | | | | IL-13 | – | – | –0.71 (–1.16 to –0.25) | **0.0026** |
| | 5 years | GM | 81 | 45 | Glutamate | Pro-inflammatory | IL-8 | –0.21 (–0.37 to –0.06) | **0.0069** | – | – |
| | | | | | Myo-inositol | Pro-inflammatory | IL-8 | –0.19 (–0.35 to –0.04) | **0.0150** | – | – |
| | | | | | | | TNFα | –0.44 (–0.71 to –0.16) | **0.0023** | – | – |
| | | | | | | Anti-inflammatory | IL-4 | –0.22 (–0.40 to –0.04) | **0.0194** | 0.22 (0.01 to 0.44) | **0.0387** |
| | | | | | | | IL-13 | –0.20 (–0.36 to –0.04) | **0.0168** | – | – |
| | | | | | | Neuroinflammatory | NGAL | –0.67 (–1.26 to –0.08) | **0.0275** | – | – |
| | | | | | Choline | Pro-inflammatory | IL-1β | – | – | –0.65 (–1.16 to –0.14) | **0.0134** |

Linear regression models with robust standard errors. Adjusted for child age, child sex, voxel composition, and **exclusive breastfeeding duration**.

### Supplementary Table 7. Early Learning Outcome Measure scores at 6–7 years

| **Developmental domain** | **CHU** (n=116) | **CHEU** (n=56) | **β** | **95% CI** | **P-value** |
|---|---|---|---|---|---|
| Cognitive development | 10.7 (5.7) | 10.7 (5.0) | 0.50 | (-0.57 to 1.93) | 0.39 |
| Fine motor coordination and visual motor integration | 17.8 (2.4) | 17.9 (3.2) | 0.00 | (-0.85 to 0.24) | 0.57 |
| Gross motor development | 11.5 (6.7) | 12.9 (7.8) | -0.03 | (-1.63 to 1.47) | 0.81 |
| Emergent mathematics | 13.6 (4.5) | 14.6 (7.1) | 0.00 | (-1.34 to 1.24) | 0.88 |
| **Emergent literacy and language** | **13.6** (**6.2**) | **11.2** (**5.3**) | **1.95** | (**0.53 to 3.37**) | **0.0084** |
| Overall school readiness score | 68.1 (15.5) | 65.7 (17.2) | 1.94 | (-1.92 to 5.93) | 0.33 |

Wilcoxon Rank Sum (Mann Whitney U) tests. Group data is median (IQR). **CHEU**: Children who are HIV-Exposed and Uninfected; **CHU**: Children who are HIV-Unexposed; **β**: Effect size. More information on the South African developmental tool *Early Learning Outcome Measures* can be found at: https://datadrive2030.co.za/elom-origins/

### Supplementary Figures

### Supplementary Figure 1. Maternal serum marker levels during pregnancy

Raw data plotted. Statistical tests were conducted on log-scaled data.

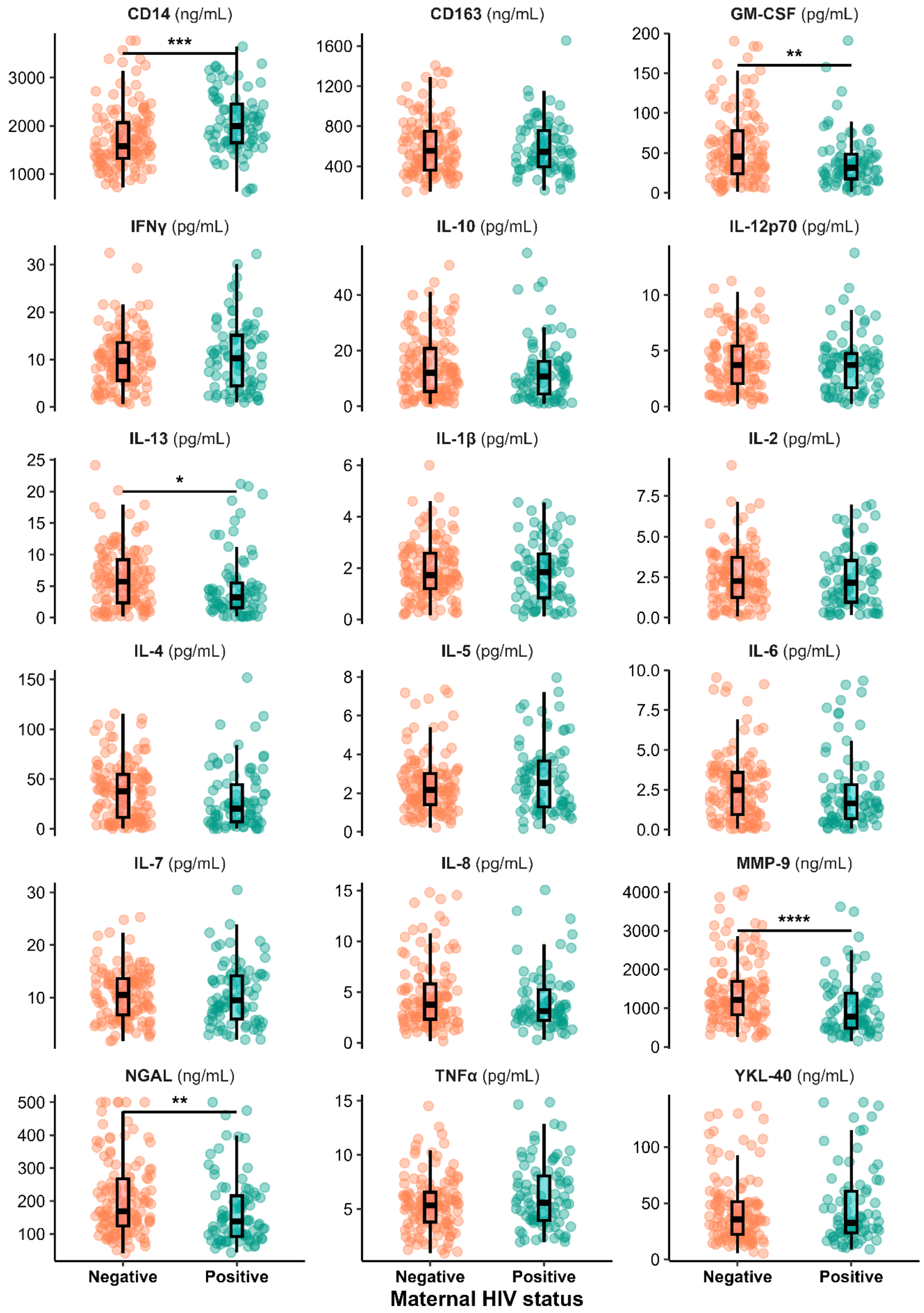

**Supplementary Figure 2. Child neurometabolite absolute concentrations at 6–7 years of age**

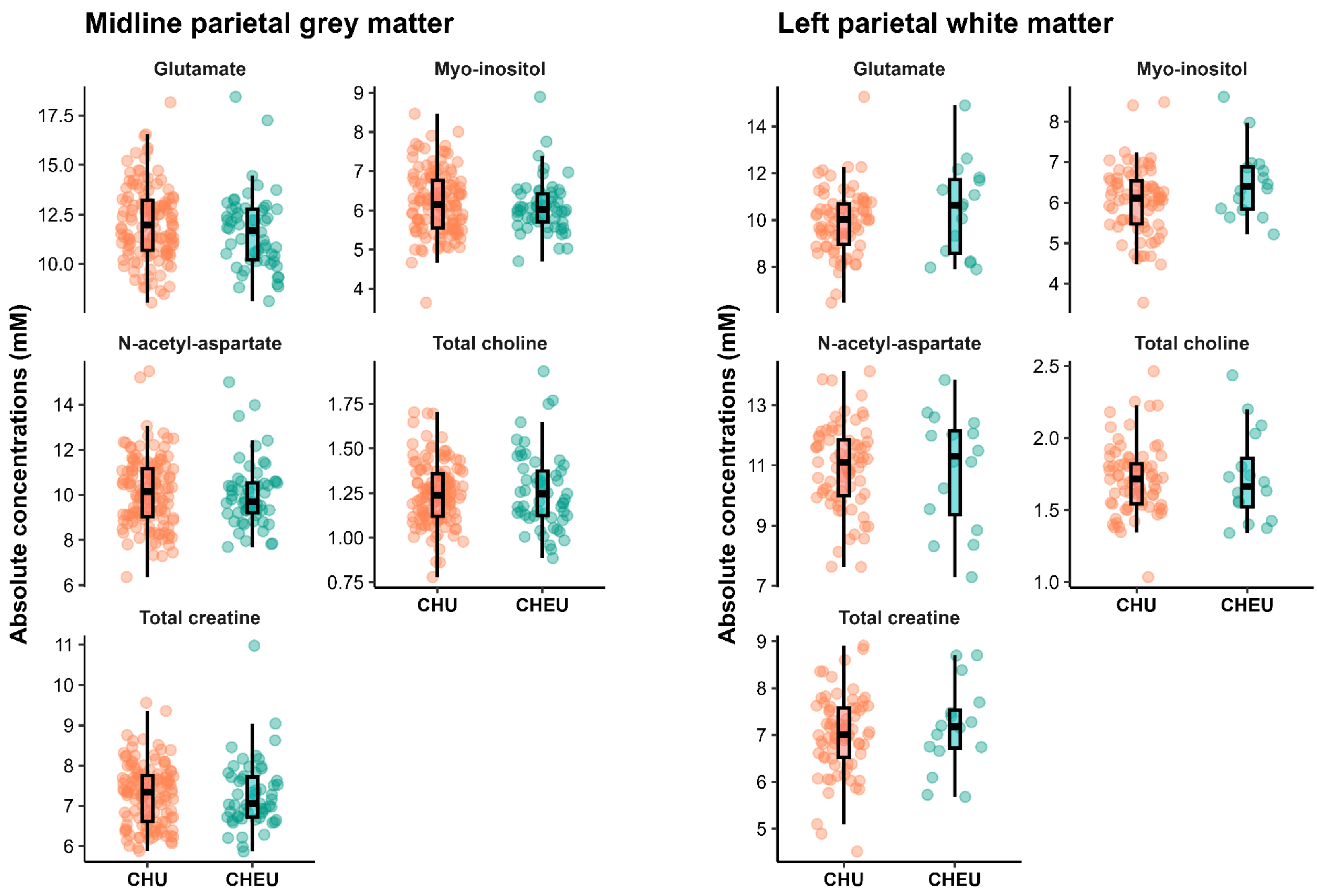

### Mediation analyses

To investigate whether maternal/child serum immune markers and/or child neurometabolite ratios mediated the relationship between maternal HIV status and child neurodevelopmental outcomes, three sets of mediation models were conducted using structural equation modeling (SEM) within the *lavaan* package in R. All models were fitted using full information maximum likelihood (FIML) to account for missing data and non-parametric bootstrapping with 5,000 resamples to obtain robust standard errors and percentile-based 95% confidence intervals for indirect effects.

Model specifications were guided by prior biological hypotheses and upstream analyses that identified significant associations between maternal HIV status, maternal/child serum markers, child neurometabolite ratios, and Early Learning Outcomes Measure (ELOM) scores at 6–7 years of age. Separate models were run for each candidate serum marker or marker slope, neurometabolite ratio, or marker–neurometabolite pair.

### Model 1

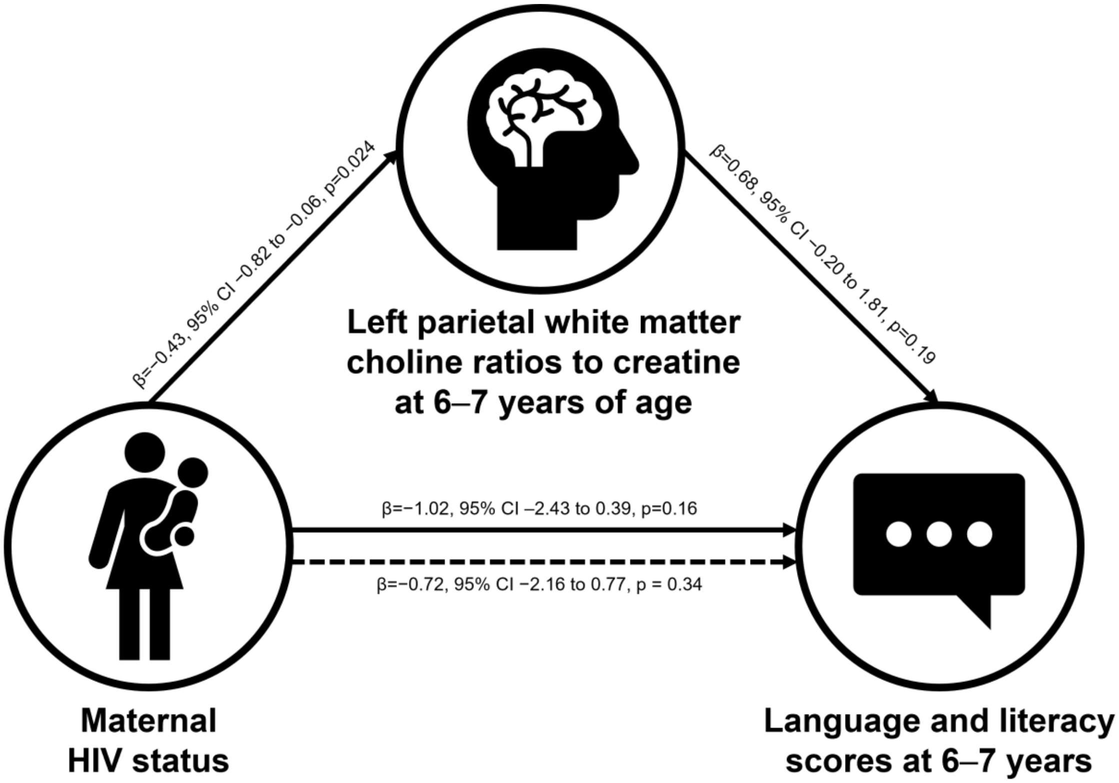

The first model evaluated whether left parietal white matter choline ratios to creatine at 6–7 years mediated the association between maternal HIV status and child ELOM language and literacy scores. In group comparisons, both choline ratios (**Supplementary Table 1**) and ELOM language and literacy scores (**Supplementary Table 6**) were significantly lower in children who are HEU compared to HU peers. Within the HU group, higher left parietal white matter choline ratios were positively associated with better language and literacy performance. These patterns provided the biological and statistical rationale for testing whether differences in choline levels could explain, at least in part, the relationship between maternal HIV exposure and child neurodevelopmental outcomes.

In this model, maternal HIV was significantly associated with lower choline ratios (β=−0.43, 95% CI −0.82 to −0.06, p=0.024). Higher choline ratios were positively associated with ELOM language and literacy scores (β=0.68, 95% CI −0.20 to 1.81, p=0.19), although this association was not statistically significant. The indirect effect of maternal HIV on ELOM scores through choline was negative but non-significant (β=−0.30, 95% CI −0.91 to 0.12, p=0.26). The total effect of maternal HIV on language and literacy scores was β=−1.02, 95% CI –2.43 to 0.39, p=0.16, and the direct effect, after accounting for choline, was β=−0.72, 95% CI −2.16 to 0.77, p = 0.34.

#### Model 2

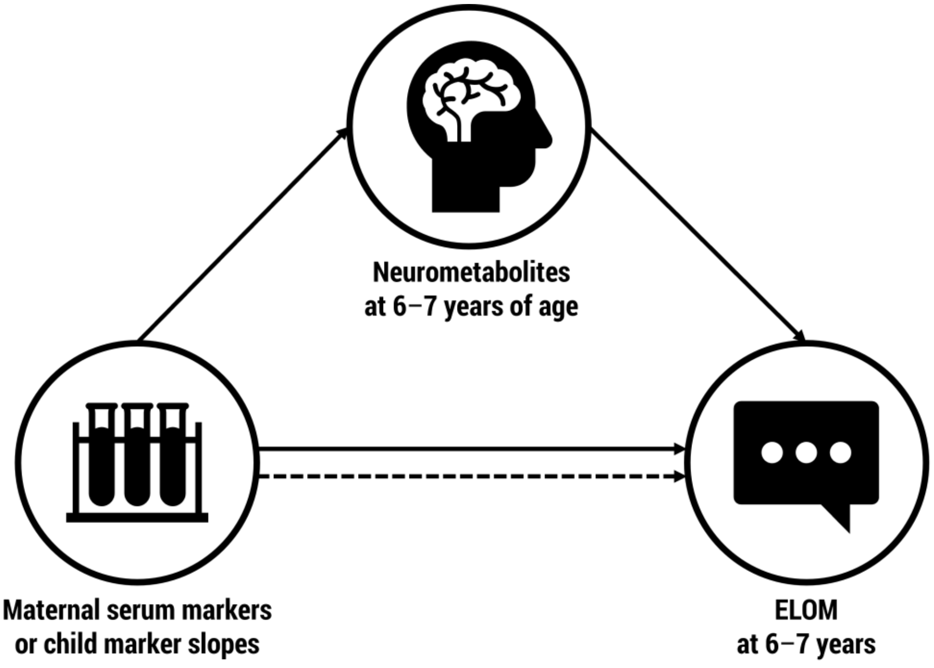

The second set of analyses extended this framework to systematically evaluate whether selected serum markers, neurometabolite ratios in either midline parietal grey matter or left parietal white matter, or marker–metabolite pairs acted as mediators of the maternal HIV → child ELOM pathway.

For each model, a different candidate mediator was explored based on upstream associations (**Table 2** and **Table 3** in main manuscript). The model structure remained the same as in Model 1, with maternal HIV as the predictor and child ELOM scores as the outcome. Each mediation model was run independently, and none reached our threshold for statistical significance.

### Model 3

The third set of analyses tested serial multiple mediation to capture a potential biological cascade where maternal or child serum markers influence child neurometabolite levels, which in turn affect neurodevelopment and school readiness.

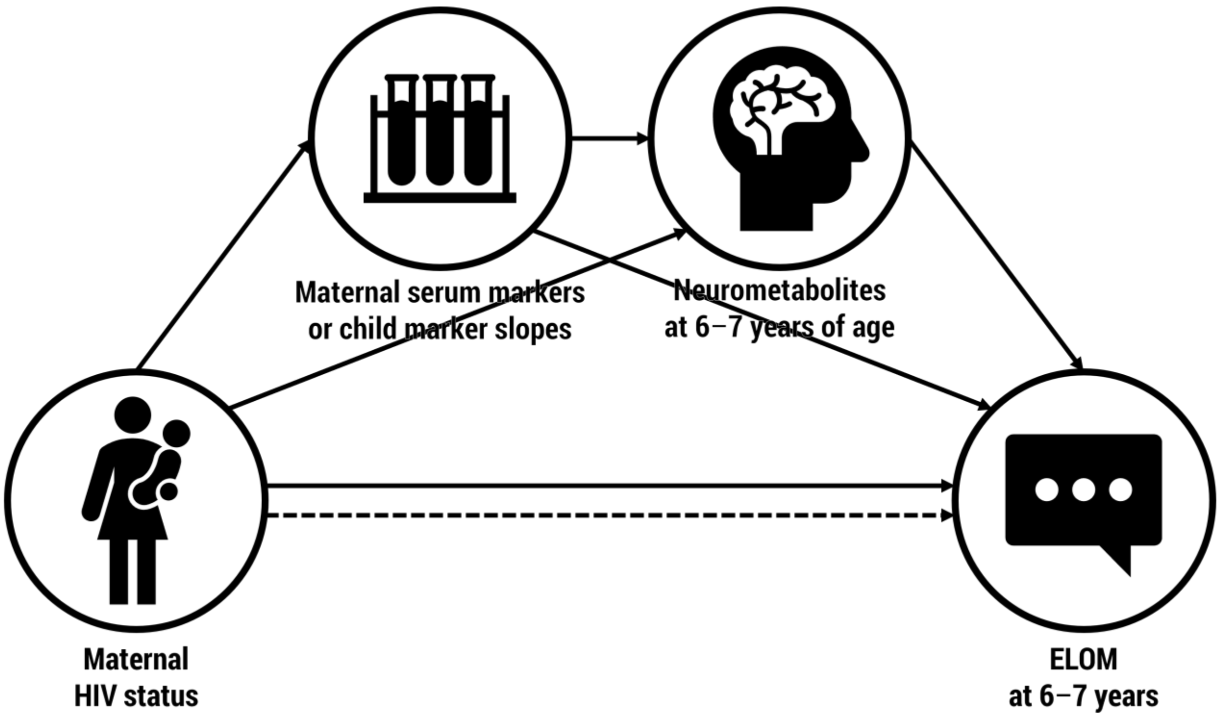

This model included two mediators in sequence:

- Predictor (X): Maternal HIV status
- Mediator 1 (M1): Serum marker (maternal or child, cross-sectional or longitudinal)

- Mediator 2 (M2): Child neurometabolite ratios to creatine (parietal grey or white matter)
- Outcome (Y): ELOM score (overall or domain-specific)

The indirect effects were decomposed into three components:

- X → M1 → Y
- X → M2 → Y
- X → M1 → M2 → Y (serial pathway)

The total indirect effect was calculated as the sum of these three pathways. Parameter estimates, standardized coefficients, and bootstrap confidence intervals were extracted to determine whether any of the indirect effects were statistically significant. Model fit indices were examined to ensure adequate model specification.

Different combinations of candidate mediators and outcomes were explored based on upstream associations. For example, IL-1β trajectories were significantly different between HEU and HU children (**Supplementary Table 3**). At age 5 years, child IL-1β concentrations were negatively associated with left parietal white matter choline ratios in the HEU group (**Table 2** in main manuscript). These choline ratios were lower in HEU compared to HU children in group comparisons (**Supplementary Table 1**), and within the HU group, higher choline ratios were positively associated with ELOM language scores. Language scores were lower in HEU versus HU children (**Supplementary Table 6**). Together, these patterns suggested a biologically plausible serial pathway in which maternal HIV status alters child early-life IL-1β trajectories, which in turn impact mid-childhood left parietal white matter choline ratios and, ultimately, language outcomes at school entry.

Each mediation model was run independently, and none reached our threshold for statistical significance.
